# Fission yeast NDR/LATS kinase Orb6 regulates exocytosis via phosphorylation of exocyst complex

**DOI:** 10.1101/291468

**Authors:** Ye Dee Tay, Marcin Leda, Christos Spanos, Juri Rappsilber, Andrew B. Goryachev, Kenneth E. Sawin

## Abstract

NDR/LATS kinases regulate multiple aspects of cell polarity and morphogenesis from yeast to mammals, but few of their substrates are known. Fission yeast NDR/LATS kinase Orb6 has been proposed to control cell polarity via spatial regulation of Gef1, a guanine nucleotide exchange factor for the small GTPase Cdc42. Here we show that Orb6 plays a critical role as a positive regulator of exocytosis, independent of Gef1. Through Orb6 inhibition *in vivo* and quantitative global phosphoproteomics, we identify several proteins involved in membrane trafficking as Orb6 targets, and we confirm Sec3 and Sec5, conserved components of the exocyst complex, as substrates of Orb6 both *in vivo* and *in vitro.* Our results suggest that Orb6 kinase activity is crucial for exocyst localization to actively-growing cell tips and for exocyst activity during septum dissolution after cytokinesis. We further show that Orb6 phosphorylation of Sec3 serine-201 contributes to exocyst function in parallel with exocyst protein Exo70. We propose that Orb6 contributes to polarized growth by regulating membrane trafficking at multiple levels.

## INTRODUCTION

The ability to establish, maintain and alter polarity is central to the function of nearly all types of eukaryotic cells^1–5^. Dysregulation of cell polarity is associated with multiple pathologies, including tumorigenesis and neurodegenerative disease^6–8^. NDR/LATS kinases (here referred to as NDR kinases) are members of a subfamily of the AGC serine/threonine kinases and are important for polarized cellular differentiation in multiple systems^9^. NDR kinases such as Trc (*Drosophila*), NDR1 and NDR2 (NDR1/2; mammals), and SAX-1/SAX-2 (*C. elegans*) share evolutionarily conserved functions in coordinating neurite branching and patterning of neuronal fields^10^. Other NDR kinases such as Wts (*Drosophila*), LATS1 and LATS2 (LATS1/2; mammals) and WTS-1 (*C. elegans*) regulate polarized differentiation of epithelia and other cell types^11^. In budding yeast *S. cerevisiae* and fission yeast *S. pombe*, NDR kinases Cbk1 and Orb6 (respectively) are required for polarized cellular morphogenesis^12–14^. Other essential biological processes regulated by NDR kinases include centrosome duplication, cell cycle progression, autophagy and apoptosis^10^.

Identification of targets of NDR kinases is crucial for understanding how they regulate complex biological processes. To date, surprisingly few targets of NDR kinases are known^15^. The best characterized targets in mammals and *Drosophila* are the transcriptional co-activators YAP and TAZ (targets of LATS1/2) and Yki1 (target of Wts), respectively^16^. Phosphorylation of YAP/TAZ and Yki is an important element of the Hippo pathway, a tumour suppressor pathway regulating cell shape and proliferation^17^. In addition, NDR1/2 phosphorylate p21 cyclin-dependent kinase inhibitor and MYPT1 phosphatase, which regulate the G1/S transition and G2 DNA damage checkpoint, respectively^18, 19^. In neurons, NDR1/2 phosphorylate AP2-associated kinase 1 (AAK1) and Rab8 GTPase guanine nucleotide-exchange factor (GEF) Rabin8, which are involved in vesicle trafficking and are important for dendrite growth regulation and dendritic spine development, respectively^20^.

In budding yeast, inactivation or inhibition of Cbk1 affects both cell morphogenesis and asymmetry of gene expression between mother and daughter cell. Cbk1 phosphorylates the transcription factor Ace2 and the RNA-binding protein Ssd1, a translational regulator^21^. Cbk1 is also reported to phosphorylate Sec2, a GEF for the Rab GTPase Sec4^22^. In fission yeast, temperature-sensitive *orb6-25* mutants lose polarity at non-permissive temperature, and cells become round rather than rod-shaped^12^. *In vitro*, Orb6 can phosphorylate the Ssd1 homolog Sts5^23^, as well as Gef1, a GEF for the Rho-family cell-polarity GTPase Cdc42^24^. Phosphorylation of Gef1 is thought to promote its association with 14-3-3 protein Rad24, restricting Gef1’s ability to activate Cdc42^24^. Accordingly, Orb6 inactivation leads to ectopic localization of active Cdc42 (Cdc42-GTP) on cell sides, with subsequent recruitment of formin For3 to these ectopic sites^25^. These events have been proposed to drive reorganization of actin cable nucleation and redirection of intracellular transport, thus leading to increased cell width in Orb6-inhibited cells^25^.

Here we show, in contrast to previous work, that several phenotypes associated with Orb6 inactivation, including increased cell width, are independent of Gef1. We further find that Orb6 positively regulates exocytosis, also independently of Gef1. To identify novel targets of Orb6, we performed a quantitative global phosphoproteomics analyses of cells with decreased Orb6 kinase activity and generated a high-quality dataset of Orb6 targets *in vivo*. Targets include proteins involved in kinase signalling, membrane trafficking, and the exocyst complex, a conserved octameric complex involved in vesicle tethering prior to membrane fusion^26–28^. We further show that Orb6 phosphorylates exocyst *in vitro* and that exocyst phosphorylation *in vivo* mediates a subset of Orb6-dependent phenotypes. Overall, our results suggest that Orb6 regulates cell polarity via multiple targets involved in membrane trafficking and exocytosis.

## RESULTS

### Several cell growth/polarity phenotypes after Orb6 inhibition in vivo are Gef1-independent

A powerful way to study protein kinase function *in vivo* involves mutating “gatekeeper” residues in the ATP-binding pocket, making kinase activity sensitive to cell-permeable nucleotide-competitive analogs^29^. An earlier analog-sensitive *orb6* allele, *orb6-as2*, was driven by the high-strength *nmt1* promoter^25^. To investigate Orb6 kinase function at native expression levels, we introduced the same mutation (M170A) at the endogenous *orb6* locus; throughout this work we will refer to this new allele as *orb6-as2.* We inhibited Orb6-as2 kinase activity using the nucleotide competitive analog 3-BrB-PP1, although other analogs were equally effective (Suppl. Fig. 1A,B). For simplicity, we will refer to our ***orb6-as2*** cells treated with 3-BrB-PP1 as “Orb6-inhibited” cells. Consistent with previous work^23, 25^, Orb6 inhibition *in vivo* led to an increase in cell width, as well as some ectopic localization of Cdc42-GTP and the actin cytoskeleton to cell sides, and a defect in cell separation during septation (Fig. 1A,C; Suppl. Movie 1). Also consistent with previous work, ectopic Cdc42-GTP and actin localization upon Orb6 inhibition was suppressed by *gef1∆* (Suppl. Movie 1)^25^.

**Figure 1.**
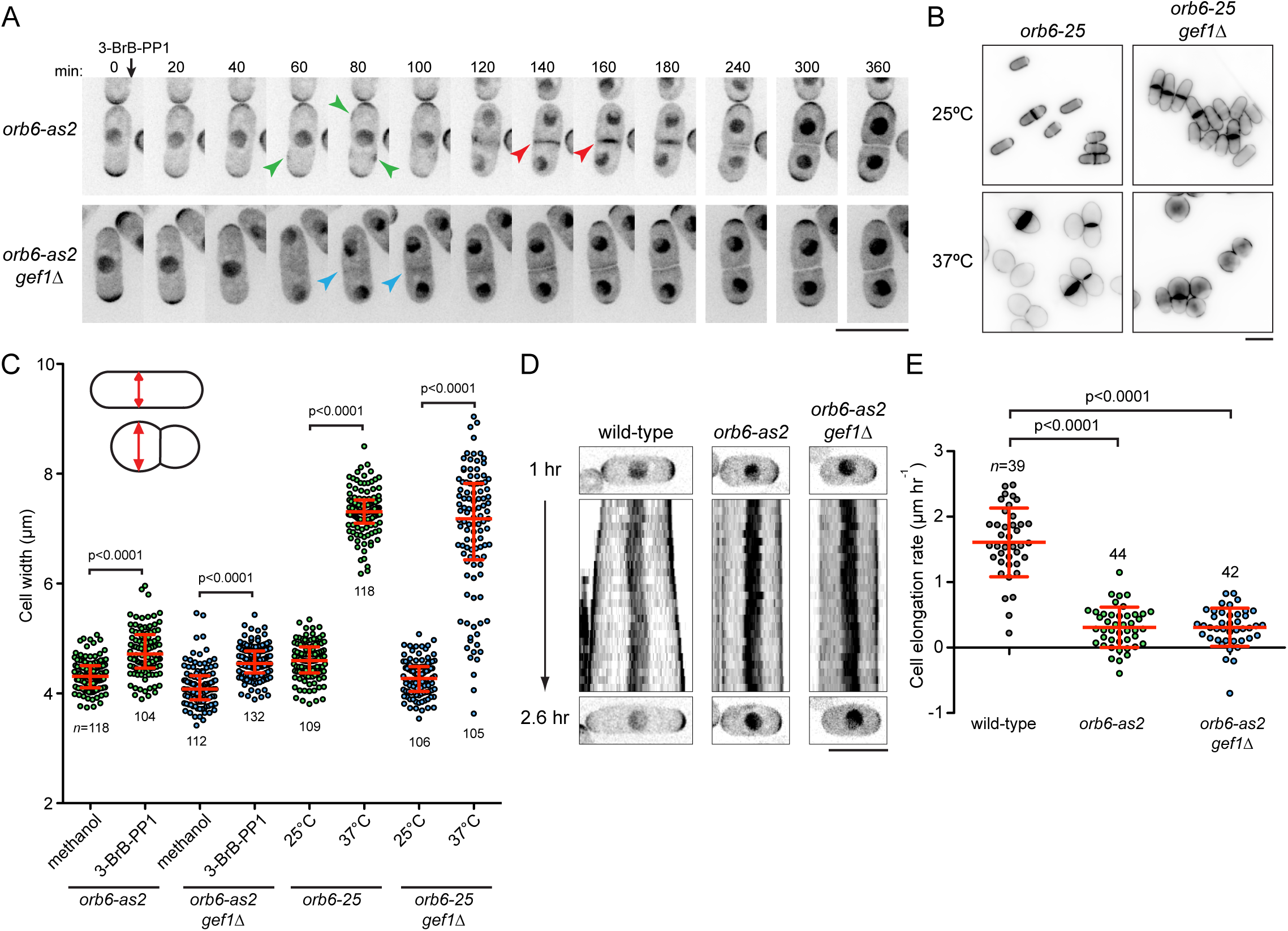
Orb6 inhibition leads to increased cell width, cell-separation defects, and cessation of polarized elongation in both wild-type and gef1∆cells. **(A)** Timepoints from movies of Cdc42-GTP reporter CRIB-3mCitrine in indicated strains upon Orb6 inhibition. 3-BrB-PP1 was added after zero timepoint. Larger fields including these cells (and also showing Lifeact-3mCherry) are shown in Supplementary Movie 1. Green arrows indicate ectopic CRIB-m3Citrine on cell sides in *orb6-as2* cells but not *orb6-as2 gef1∆* cells. Red arrows indicate CRIB-m3Citrine at midzone during early stages of septation in *orb6-as2* cells. Blue arrows indicate absence of CRIB-m3Citrine at comparable stages in *orb6-as2 gef1∆* cells (see Supplementary Movie 1). Both strains show width increase and cell-separation defects. **(B)** Calcofluor images of indicated strains at 25°C and after 5 hr at 37°C. **(C)** Cell width in the strains and conditions indicated, after 5 hr treatment. Diagrams illustrate width measurement. Red lines and error bars show median and interquartile range. *n* indicates number of cells scored. **(D)** Kymographs of CRIB-3mCitrine in indicated strains, starting 1 hr after 3-BrB-PP1 addition. The same cells are shown in Supplementary Movie 2. Orb6-inhibited cells do not elongate, despite enrichment of Cdc42-GTP at cell tips. **(E)** Cell elongation rates from experiments as shown in (D). Red line and error bars show mean and SD. Inspection of movies shows that slight positive values in *orb6-as2* and *orb6-as2 gef1∆* cells are due to swelling of cells in all directions and not to polarized elongation (see Supplementary Movie 2). Bars, 10 µm.

However, in apparent contrast to earlier work^25^, we found that increased cell width in Orb6-inhibited cells was not suppressed by *gef1∆* (Fig. 1A,C; Suppl. Movie 1). Because earlier work investigated *gef1∆* suppression of cell-width increase only in temperature-sensitive *orb6-25* mutants and not in Orb6-inhibited cells, we also examined *orb6-25* mutants. In liquid culture, the increase in cell width in *orb6-25* mutants (at restrictive temperature) is much greater than in Orb6-inhibited cells, such that many *orb6-25* cells become completely round (Fig. 1B,C)^12^. In any case, similar to Orb6-inhibited cells, we found that the cell-width increase in *orb6-25* cells was not suppressed by *gef1∆*, although a small proportion of cells (~16%) displayed widths similar to *orb6-25* cells at permissive temperature (Fig. 1B,C).

Interestingly, time-lapse observation of Cdc42-GTP localization revealed that Orb6-inhibited cells almost completely ceased to elongate even when they remained largely polarized (based on retention of Cdc42-GTP at cell tips; Fig. 1D,E; Suppl. Movie 2). This was surprising because Cdc42-GTP enrichment on the plasma membrane is normally closely correlated with active cell growth (Suppl. Fig. 1C)^30^. Importantly, cessation of elongation also occurred in Orb6-inhibited *gef1∆* cells, which do not show any ectopic Cdc42-GTP localization.

We conclude that in addition to increased cell width and impaired cell separation, a key phenotype associated with Orb6 inhibition *in vivo* is the cessation of cell elongation, independent of changes in Cdc42-GTP at cell tips. Moreover, none of these phenotypes is suppressed by *gef1∆*.

### Orb6 inhibition leads to a strong decrease in exocytosis

To further characterize cessation of elongation in Orb6-inhibited cells, we imaged mCherry-tagged beta-glucan synthase Bgs4, a multipass transmembrane protein required for cell wall synthesis at zones of active growth (i.e. the cell tips during interphase, and the cell midzone during cytokinesis^31, 32^ (Suppl. Fig. 1C). Within 12-15 minutes after Orb6 inhibition, Bgs4 was almost completely lost from cell tips, in both wild-type (i.e. *gef1+*) and *gef1∆* genetic backgrounds (Fig. 2A,B,D; Suppl. Movie 3). Loss of Bgs4 from cell tips was accompanied by an increase in cytoplasmic Bgs4 puncta, suggesting that Bgs4 was in endomembrane compartments. By contrast, control wild-type (*orb6^+^*) cells treated with 3-BrB-PP1 showed no change in mCherry-Bgs4 localization and continued to elongate (Fig. 2A; Suppl. Fig. 1D).

**Figure 2.**
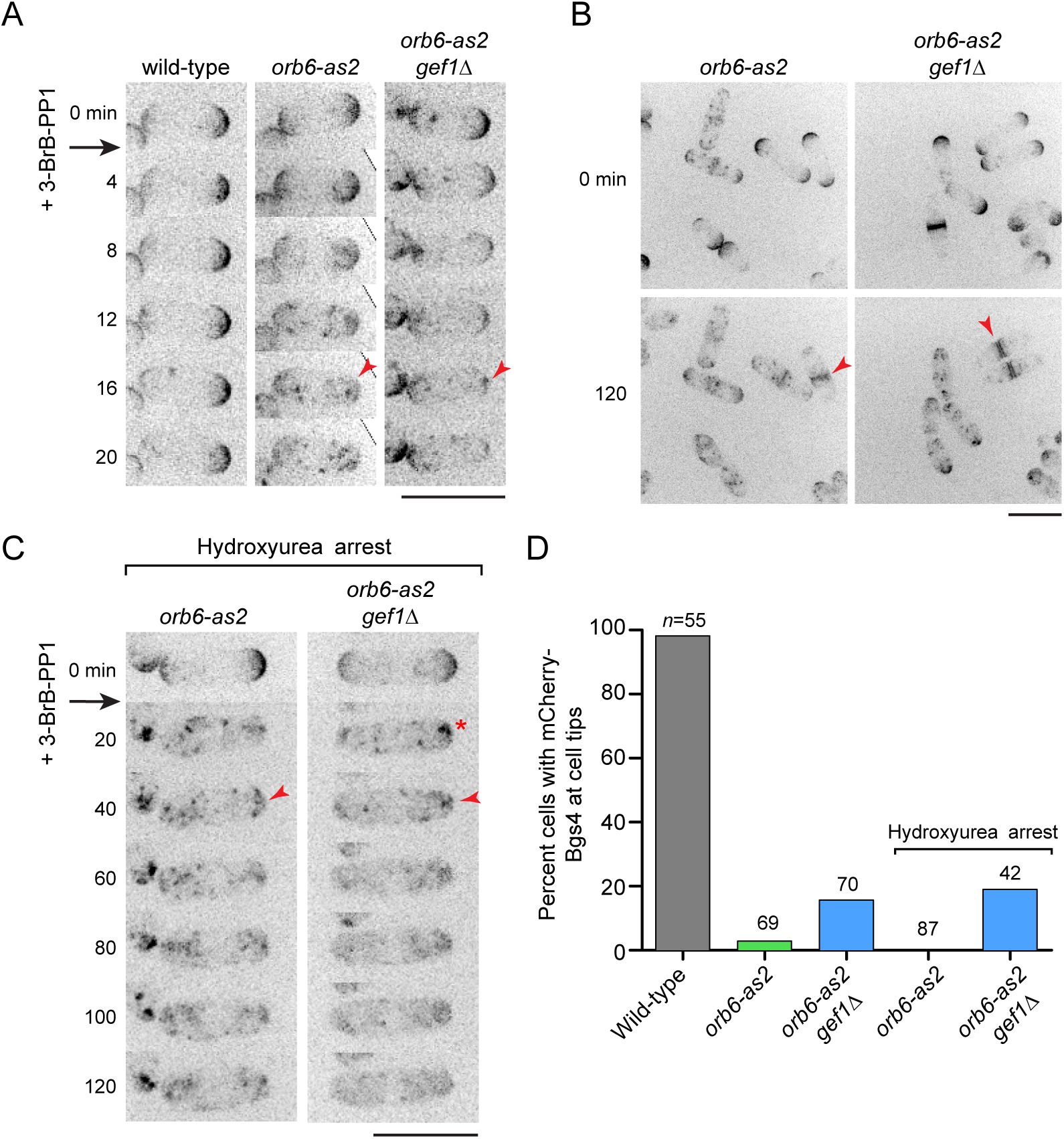
After Orb6 inhibition, integral membrane protein Bgs4 (beta-glucan synthase) is lost from the plasma membrane at interphase cell tips, in both wild-type and gef1∆ cells. **(A)** Timepoints from movies showing mCherry-Bgs4 after Orb6 inhibition in the indicated strains. 3-BrB-PP1 was added after zero time point. Arrowheads indicate loss of Bgs4 from cell tips. **(B)** Matched fields of cells before (0 min) and after (120 min) 3-BrB-PP1 addition. Arrowheads indicate accumulation of Bgs4 at septum. The same cells are shown in Supplementary Movie 3. **(C)** Timepoints from movies showing mCherry-Bgs4 after Orb6 inhibition in hydroxyurea-arrested (G1/S) cells. Arrowheads indicate sustained loss of Bgs4 from cell tips, independent of karyokinesis. Asterisk indicates cluster of endocytosed Bgs4, under the plasma membrane. Larger fields including these cells are shown in Supplementary Movie 3. **(D)** Quantification of mCherry-Bgs4 cell-tip localization, 120 min after 3-BrB-PP1addition. *n* indicates number of cells scored. Bars, 10 µm.

Consistent with cell-separation defects, in Orb6-inhibited cells attempting to divide, Bgs4 accumulated at the septation zone for long periods (Fig. 2B; Suppl. Movie 3). Because nearly all Orb6-inhibited interphase cells eventually progressed into mitosis, we wanted to confirm that loss of Bgs4 from cell tips after Orb6 inhibition during interphase was not necessarily associated with subsequent recruitment of Bgs4 to the septation zone during division. We therefore arrested *orb6-as2* cells in G1/S with hydroxyurea and then inhibited Orb6. In the arrested cells, Bgs4 was quickly lost from cell tips, for the duration of imaging (~2 hours), and we observed similar results in Orb6-inhibited *gef1∆* cells (Fig. 2C,D; Suppl. Movie 3). Collectively, these results suggest that Orb6 kinase activity is essential for maintaining plasma membrane localization of Bgs4, and hence cell elongation, at interphase cell tips.

Because Bgs4 is a multipass transmembrane protein, its disappearance from cell tips after Orb6 inhibition should be mediated by endocytosis. To confirm this, we blocked endocytosis in *orb6-as2* cells by depolymerizing the actin cytoskeleton with Latrunculin A (LatA)^33^, and we then inhibited Orb6. In these cells, Bgs4 remained at cell tips for at least 2 hours after Orb6 inhibition (Fig. 3A; Suppl. Movie 4), strongly suggesting that loss of Bgs4 from cell tips after Orb6 inhibition occurs via endocytosis.

**Figure 3.**
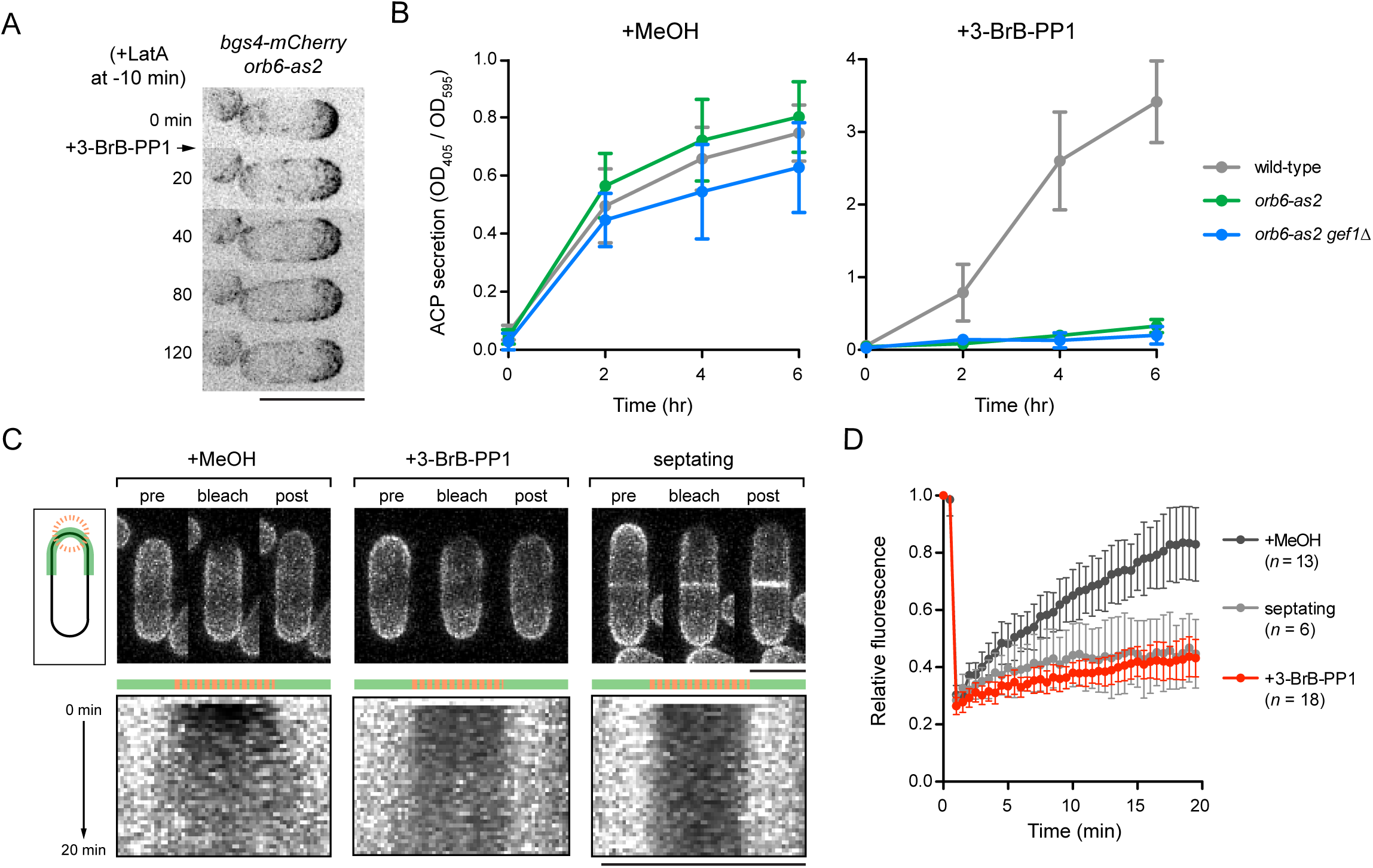
Exocytosis defects in wild-type and *gef1∆* cells after Orb6 inhibition. **(A)** Timepoints from movies showing mCherry-Bgs4 after combined actin depolymerization by Latrunculin A (LatA) and Orb6 inhibition. Disruption of endocytosis by LatA prevents loss of Bgs4 from cell tips after Orb6 inhibition (compare with Fig. 2A). A larger field including this cell is shown in Supplementary Movie 4, together with a field of *gef1∆* cells after actin depolymerization and Orb6 inhibition. **(B)** Acid phosphatase (ACP) secretion into culture medium in the indicated strains after treatment with methanol (left) or 30µM 3-BrB-PP1 (right). Orb6-inhibited cells are defective in ACP secretion. Graphs show mean ACP activity normalized to cell density, from replicate experiments. Error bars indicate SEM. **(C)** Fluorescence recovery after photobleaching (FRAP) of plasma membrane t-SNARE GFP-Psy1 at cell tips in *orb6-as2* cells under the conditions indicated. Top images are representative experiments. Bottom images are corresponding kymographs based on the line (green) and bleached region (orange) shown in diagram. (D) Quantification of FRAP from experiments as in (C). Error bars indicate standard deviation. *n* indicates number of cells analyzed. Because Psy1 is a transmembrane protein that turns over via endo/exocytosis, lack of recovery after bleaching indicates impaired exocytosis. Bars, 10 µm.

Based on these results, we hypothesized that loss of Bgs4 from cell tips after Orb6 inhibition may be due to impaired exocytosis, in the presence of continued endocytosis. This could also explain, at least in part, both the cessation of cell elongation during interphase and cell-separation defects during cell division (due to inability to dissolve the primary septum;^34, 35^). To test this, we measured secretion of acid phosphatase (ACP)^34^, into the culture medium after Orb6 inhibition. In control *orb6-as2* and *orb6-as2 gef1∆* cells, ACP secretion was similar to wild-type cells (Fig. 3B). By contrast, Orb6-inhibited cells (both *gef1+* and gef1∆) showed a sharp decrease in ACP secretion. (Fig. 3B). These results suggest that Orb6 positively regulates exocytosis and that this regulation is independent of Gef1.

In these experiments we noticed that 3-BrB-PP1-treated wild-type cells showed much higher (~4-fold) ACP secretion than control wild-type cells (Fig. 3B). In mass spectrometry experiments associated with phosphoproteomics analysis after Orb6 inhibition (see below), we found that 3-BrB-PP1 treatment leads to increased intracellular levels of the ACP Pho1 (Suppl. Fig. 2B). This likely accounts for the increase in ACP secretion seen in 3-BrB-PP1-treated vs. control cells. Importantly, 3-BrB-PP1 treatment also led to increased levels of Pho1 in *orb6-as2* cells (Suppl. Fig. 2B). These results confirm that the decrease in ACP secretion observed in Orb6-inhibited cells is due to a decrease in exocytosis and not simply to a decrease in ACP levels relative to uninhibited cells.

To further investigate the role of Orb6 in exocytosis, we measured fluorescence recovery after photobleaching (FRAP) of the plasma membrane t-SNARE Psy1 in Orb6-inhibited cells (Fig. 3C,D). Psy1 is a singlepass transmembrane protein that is distributed approximately uniformly on the plasma membrane^36, 37^, and previous FRAP experiments showed that GFP-Psy1 recovers fluorescence when bleached at cell tips but not when bleached at cell sides^37^. This implies that Psy1 is not highly mobile in the plane of the plasma membrane and that Psy1 fluorescence recovery at cell tips is due primarily to turnover via endo- and exocytosis. Consistent with previous work, we found that GFP-Psy1 bleached at cell tips in control interphase cells recovered to approximately 80% of prebleach levels within 20 mins after bleaching, with essentially uniform recovery across the bleached zone. By contrast, GFP-Psy1 bleached at cell tips after Orb6 inhibition showed very little recovery. As an additional control, we bleached GFP-Psy1 at cell tips in control cells during septation, when membrane trafficking is concentrated at the cell midzone rather than cell tips; in these cells, we also observed little fluorescence recovery. Collectively, these results suggest that decreased fluorescence recovery of GFP-Psy1 at cell tips in Orb6-inhibited cells is due to defects in exocytic membrane trafficking.

We conclude that inhibition of Orb6 kinase activity impairs exocytosis, independently of Gef1.

### Identification of Orb6 substrates by quantitative global phosphoproteomics

To date, the only reported substrates of Orb6 are Gef1 and Sts5, both of which were phosphorylated *in vitro* by immunoprecipitates of the Orb6 coactivator Mob2^23, 24^. Sts5 is not known to be associated with exocytosis, and our results indicate that decreased exocytosis after Orb6 inhibition does not involve Gef1. This suggested that other, yet unknown, substrates of Orb6 may regulate exocytosis.

To identify novel Orb6 substrates, we combined Stable Isotope Labeling with Amino Acids in Culture (SILAC)^38–40^ with global phosphoproteomics analysis (Suppl. Fig. 2A). We treated “light” and “heavy”-labeled *orb6-as2* cells with either methanol (control) or 3-BrB-PP1 and quantitatively analyzed phosphopeptides by mass spectrometry to identify phosphosites with decreased phosphorylation after Orb6 inhibition (see Methods). Phosphopeptide abundance was normalized to the abundance of the relevant individual proteins, and to control for any possible off-target effects of 3-BrB-PP1, we also analyzed phosphopeptides in wild-type cells treated with 3-BrB-PP1.

From three biological replicate experiments we identified a total of 10,866 phosphosites, of which 8,134 could be quantified. Among quantified phosphosites, 326 (4%) showed a two-fold or greater decrease in phosphorylation after Orb6 inhibition in at least one experiment, 121 (1.5%) in at least two experiments, and 55 (0.7%) in all three experiments (Suppl. Table 1). Correlation between two representative replicate experiments is shown in Fig. 4A.

**Figure 4.**
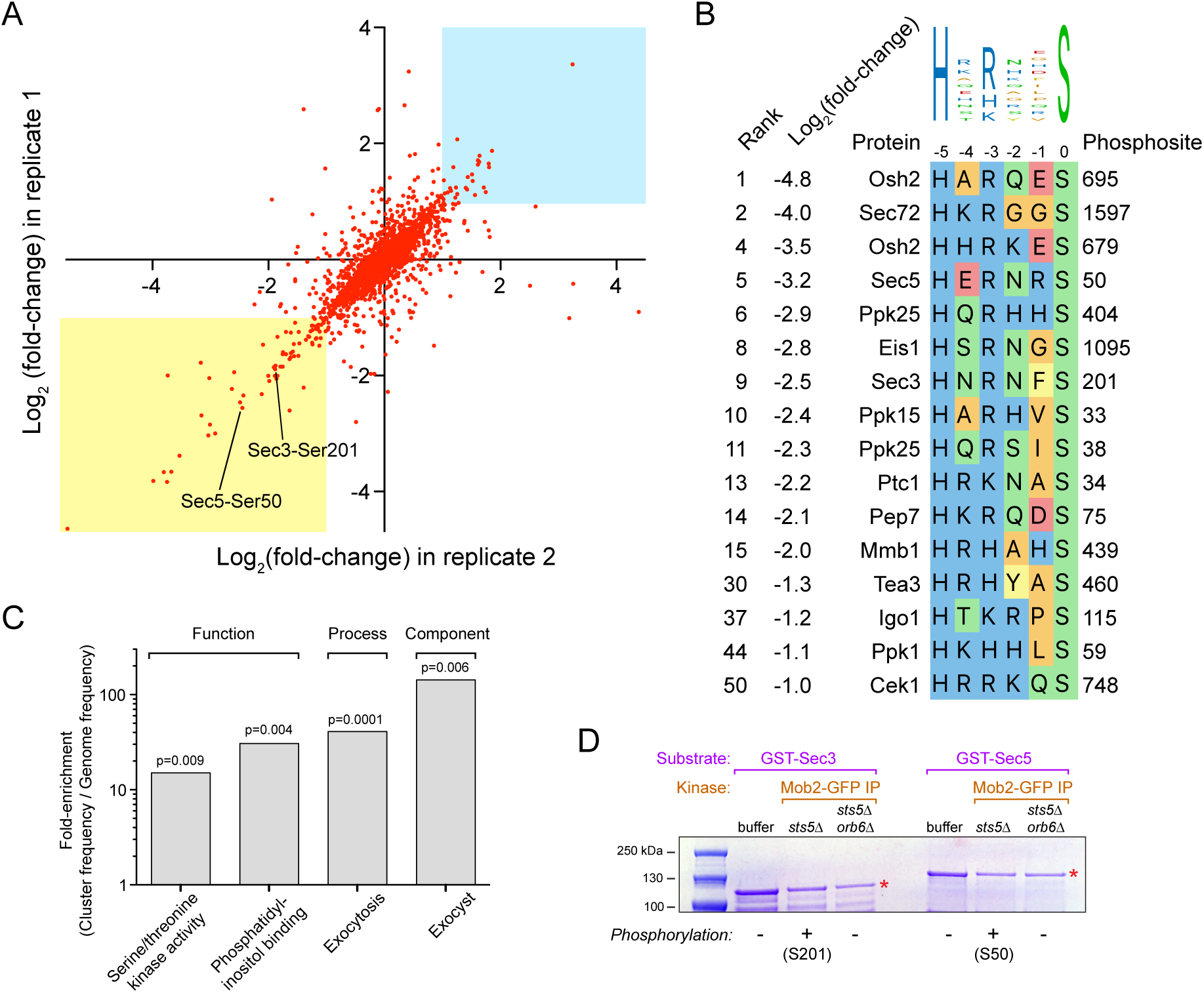
Identification of Orb6 targets *in vivo by* quantitative global phosphoproteomics. **(A)** Correlation of changes in phosphorylation after Orb6 inhibition for 2,415 quantified phosphosites common to two biological replicate experiments. Axes show log_2_(fold-change) in phosphorylation of individual phosphosites. Yellow region indicates phosphosites with two-fold or greater decreased phosphorylation after Orb6 inhibition in both experiments. Blue region indicates relative scarcity of phosphosites with an equivalent increase in phosphorylation. Phosphosites Sec3 serine-201 (S201) and Sec5 serine-50 (S50) are indicated. **(B)** The highest-confidence and highest-ranking Orb6-dependent phosphosites match the NDR/LATS consensus (HxR/H/KxxS). All phosphosites indicated were quantified in all three replicate experiments and showed two-fold or greater decreased phosphorylation after Orb6 inhibition. Phosphosites were ranked by fold-decrease in phosphorylation (i.e., independent of whether or not they matched the NDR/LATS motif). See also Supplementary Table 1. **(C)** Enrichment of gene ontology terms associated with the genes shown in (B), based on molecular function, biological process and cellular component ontologies. **(D)** Phosphorylation of Sec3 serine-201 (S201) and Sec5 serine-50 (S50)by Orb6 *in vitro*. Recombinant GST-Sec3 or GST-Sec5 (red asterisks) was incubated with buffer alone, or with Mob2-GFP immunoprecipitates from *sts5∆* or *sts5∆ orb6∆* cells. Phosphorylation on S201 and S50 was detected only in kinase reactions from Mob2-GFP IPs in *sts5∆* strains. MS phosphorylation data from the experiment is shown in Supplementary Figure 4.

Phosphosites with decreased phosphorylation after Orb6 inhibition could be either direct or indirect targets of Orb6. To classify phosphosites, we compared them to the NDR/LATS consensus motif, HxR/H/KxxS/T^41–43^. Among the 55 phosphosites with significantly decreased phosphorylation in all three experiments, 36 (65%) contained a basic amino acid (histidine, arginine or lysine) at the −3 position (Fig 4B; Suppl. Fig. 2C,D; Suppl. Table 1). NDR/LATS kinases are unique among AGC-family kinases (and possibly among all characterized serine/threonine kinases) in that they have a very strong preference for histidine at position −5 relative to the phosphorylated residue^42, 44^. Strikingly, among the 15 phosphosites with the greatest decrease in phosphorylation in all three experiments (i.e. four-fold or greater), 12 showed a perfect match to the NDR/LATS consensus, including histidine at −5, suggesting that they are direct Orb6 targets (Fig. 4B; Suppl. Table 1).

Gene ontology (GO) analysis of proteins with the highest-ranking phosphosites matching the NDR/LATS consensus revealed specific enrichment for GO terms related to protein kinase signaling (Cek1, Ppk1, Ppk15, Ppk25), exocytosis and phosphatidylinositol binding (Pep7, Sec3, Sec5, Osh2), and the exocyst complex (Sec3, Sec5) (Fig. 4C). We focused further attention on exocyst proteins Sec3 and Sec5, which showed decreased phosphorylation on residues serine-201 and serine-50, respectively.

### Orb6 inhibition leads to exocyst dephosphorylation and loss from cell tips

To confirm Orb6-dependent phosphorylation of Sec3 and Sec5, we combined SILAC with Sec5 immunoprecipitation to analyze exocyst phosphorylation in control vs. Orb6-inhibited cells. In Sec5-mCherry immunoprecipitates, levels of all other exocyst proteins (Sec3, Sec6, Sec8, Sec10, Sec15, Exo70 and Exo84) were similar between control and Orb6-inhibited cells, suggesting that exocyst is largely intact in Orb6-inhibited cells (Suppl. Fig. 3A,B). Consistent with global phosphoproteomics data, in immunoprecipitates from Orb6-inhibited cells we identified decreased phosphorylation of Sec3 serine-201 and Sec5 serine-50, as well as decreased phosphorylation of a novel site, Sec8 serine-440 (Suppl. Fig. 3C). We also found that, unlike Sec5 and Sec8, Sec3 contains numerous additional phosphorylation sites that were either modestly increased or decreased in phosphorylation after Orb6 inhibition (Suppl. Fig. 3C). No other significant differences in exocyst phosphorylation were observed.

To demonstrate Orb6 phosphorylation of Sec3 and Sec5 *in vitro*, we performed kinase assays on bacterially-expressed recombinant GST-Sec3 and GST-Sec5, using Mob2-GFP immunoprecipitates (Fig. 4D & Suppl. Fig. 4). Because Orb6 is normally essential for viability, and viable *orb6∆* cells can be recovered only in an *sts5∆* background^23^, we used Mob2-GFP immunoprecipitated from *sts5∆* cells for the kinase reaction and Mob2-GFP immunoprecipitated from *sts5∆ orb6∆* cells as a negative control. Consistent with global phosphoproteomics, we identified phosphorylation of Sec3 serine-201 on GST-Sec3 and of Sec5 serine-50 on GST-Sec5 in kinase reactions prepared from *sts5∆* cells but not from *sts5∆ orb6∆* cells or buffer-only controls.

We next examined how Orb6 inhibition affects the localization of exocyst proteins *in vivo* (Fig. 5). Prior to Orb6 inhibition, fluorescent-protein fusions with Sec3, Sec5, Sec8 and Exo70 all localized to cell tips in interphase cells and to the septation zone in dividing cells. However, after Orb6 inhibition, both Sec3 and Sec5 were almost completely lost from cell tips and formed ectopic puncta on the peripheral cortex (Fig. 5A,B). Sec8 and Exo70 were partially lost from cell tips and also formed ectopic cortical puncta (Fig. 5C,D). In addition, after Orb6 inhibition, all four proteins showed prolonged localization to the septation zone, consistent with cell-separation defects. Notably, the cell-separation defect caused by Orb6 inhibition mimics the consequences of inactivating exocyst in fission yeast^34, 45, 46^.

**Figure 5.**
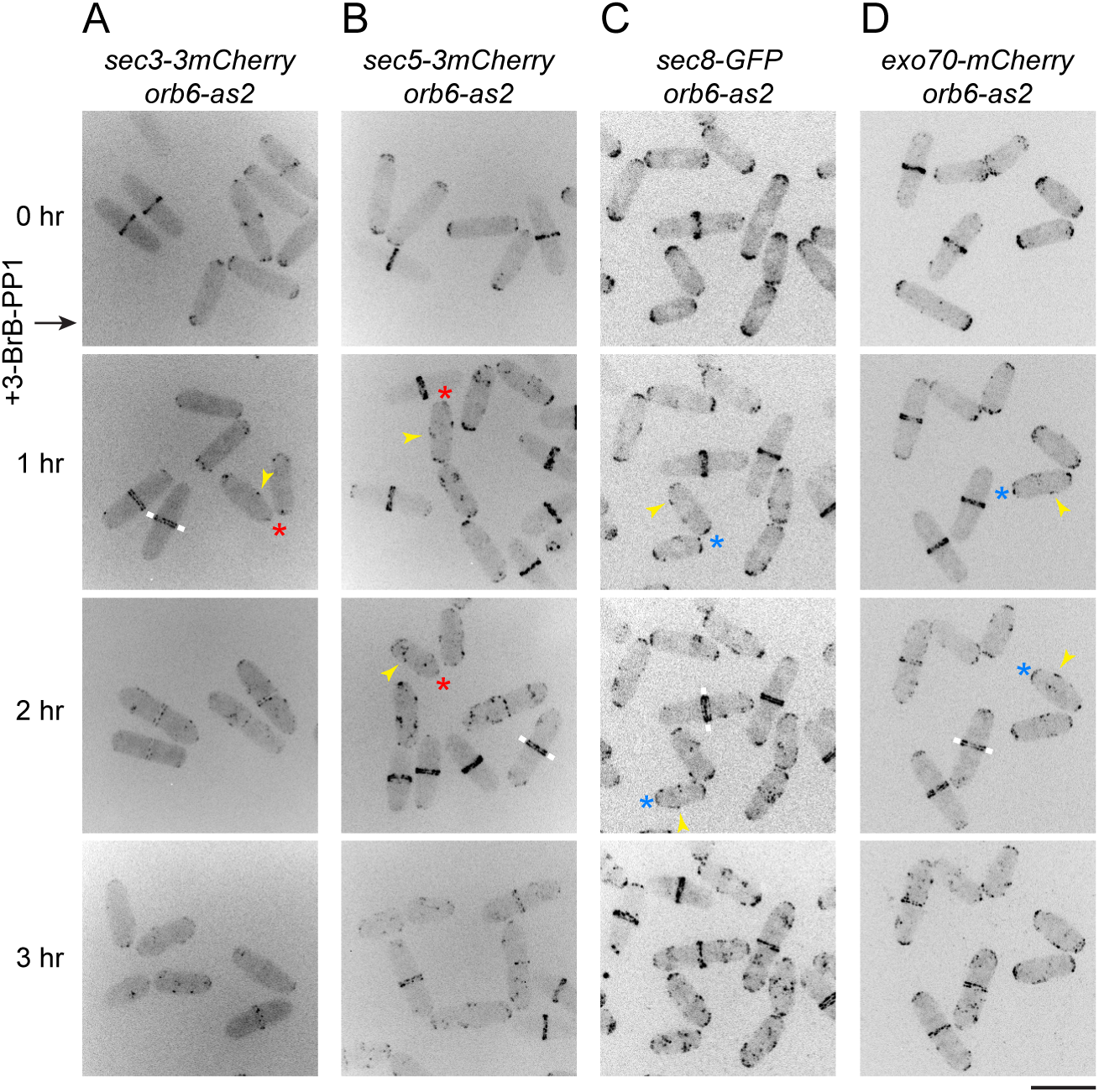
Orb6 inhibition leads to loss of exocyst from cell tips. Localization of the indicated fluorescent-protein fusions with exocyst components, before and after Orb6 inhibition. 3-BrB-PP1 was added just after the zero timepoint. Red and blue asterisks indicate complete or partial loss (respectively) of fluorescence at cell tips after Orb6 inhibition. Yellow arrowheads indicate ectopic exocyst puncta on cell sides. White bars indicate “split” exocyst localization at the septation zone. In (A) and (B), different fields were imaged for each timepoint, because of photobleaching. In (C) and (D), the same fields were imaged for all timepoints. Bar, 10 µm.

Taken together, our results indicate that Orb6 phosphorylates exocyst proteins, and Orb6 kinase activity is important for exocyst localization to the cell tip during interphase and for exocyst function in septum dissolution after cytokinesis.

### Phosphorylation of Sec3 serine-201 is important for exocyst function

To test the role of Sec3 serine-201 and Sec5 serine-50 phosphorylation in exocytosis, we mutated these residues to non-phosphorylatable alanine at the endogenous *sec3* and *sec5* loci, to make mutants *sec3-S201A* and *sec5-S50A.* In initial experiments, neither mutant displayed any obvious abnormalities similar to exocyst mutants (e.g., no cell-separation defects;^34, 45, 46^). In addition, mCherry fusions of both Sec3-S201A and Sec5-S50A localized to cell tips during interphase and to the septation zone during cell division (Fig. 6A). Given these results, we hypothesized that the function of Sec3 and/or Sec5 phosphorylation might overlap with that of other exocyst components; for example, in fission yeast, Sec3 and Exo70 are redundant for viability^45^. Consistent with this, we found that *sec3-S201A exo70∆* double mutants displayed severe cell separation defects relative to either *sec3-S201A* or *exo70∆* single mutants, with over 80% of the double-mutant cells containing at least one septum (Fig. 6B,C). Moreover, about 30% of *sec3-S201A exo70∆* cells were multi-septated. By contrast, *sec5-S50A exo70∆* double mutants displayed a septation index comparable to *sec5-S50A* and *exo70∆* single mutants, and the double mutants did not show any multi-septated phenotype (Fig. 6B,C). Consistent with these observations, *sec3-S201A exo70∆* mutants, but not *exo70∆ sec5-S50A* mutants, were strongly impaired in ACP secretion (Fig. 6D). These observations suggest that phosphorylation of Sec3 serine-201 by Orb6 works in parallel with Exo70 to promote exocyst function.

**Figure 6.**
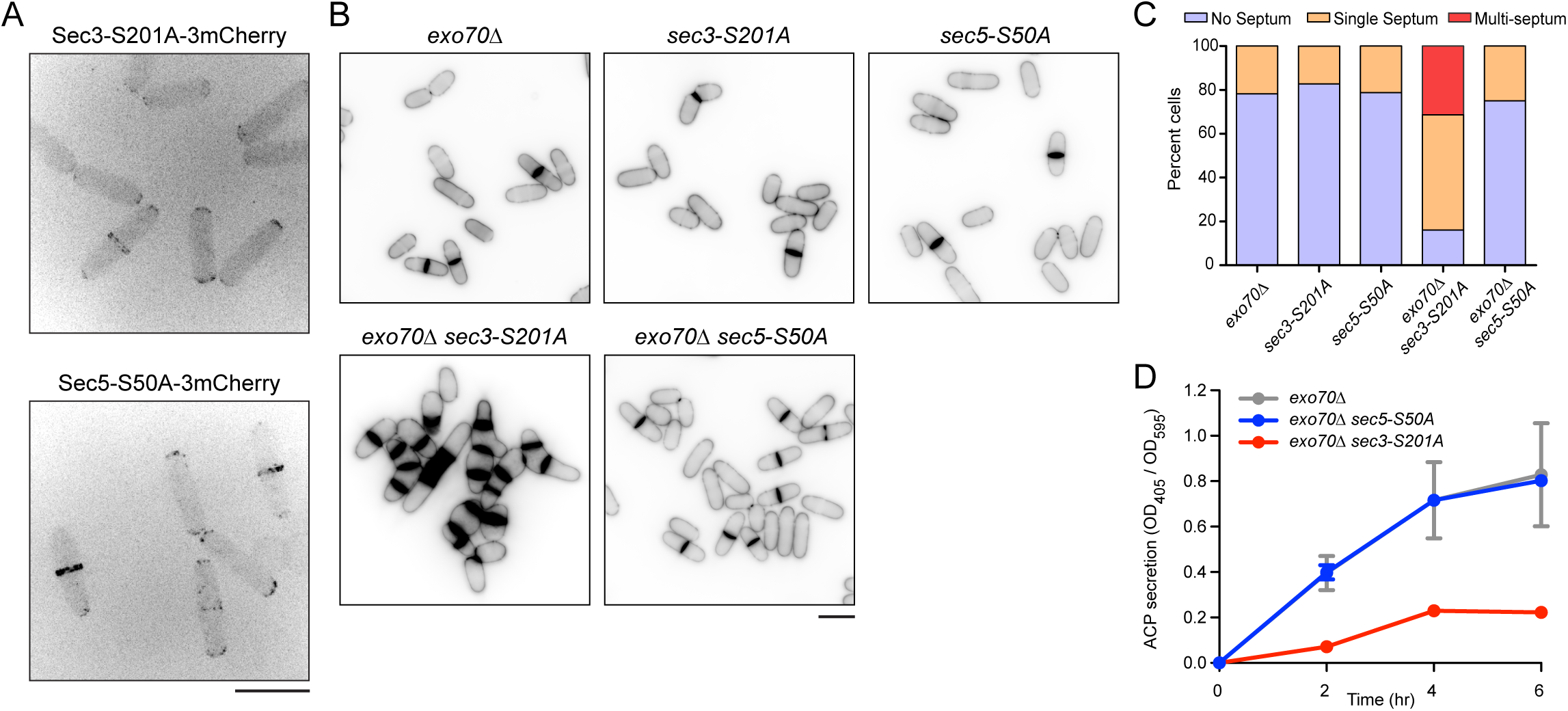
Sec3 serine-201 phosphorylation contributes to exocyst function in parallel with Exo70. **(A)** 3mCherry-tagged Sec3-S201A and Sec5-S50A mutant proteins localize normally to cell tips and septation zones. **(B)** Calcofluor staining of strains of indicated genotypes. *exo70∆ sec3-S201A* cells have strong cell-separation defects. **(C)** Quantification of septum number in cells as shown in (B). **(D)** Acid phosphatase (ACP) secretion into culture medium in the indicated strains. Graphs show mean ACP activity normalized to cell density, from replicate experiments. Error bars indicate SEM. Some error bars are smaller than the datapoint symbols and thus are not visible. Bars, 10 µm.

We hypothesized that the exocytosis and cell-separation defects in *exo70∆ sec3-S201A* cells may be due to Sec3-S201A failing to localize properly in the absence of Exo70. We attempted to examine Sec3-S201A-3mCherry localization in an *exo70∆* background, but spores of the double mutant *sec3-S201A-3mCherry exo70∆* failed to germinate (Suppl. Fig. 5A). We therefore examined Sec5 and Sec8 localization in *sec3-S201A exo70∆* cells. Both proteins localized normally to cell tips and the septation zone, suggesting that exocyst localization *per se* is not adversely affected in these cells (Suppl. Fig. 5B,C).

## DISCUSSION

Here we have shown, through a combination of microscopy, genetics and proteomics approaches, that the conserved NDR kinase Orb6 has a major role in regulating fission yeast exocytosis. Round/wide cell-polarity defects in *orb6* mutants were previously attributed to failure to spatially restrict activity of Cdc42 GEF Gef1^25^, but we find that these and other *orb6* phenotypes persist in *gef1∆* cells. Our experiments indicate that impaired exocytosis is a key Gef1-independent phenotype after inhibition of Orb6 kinase activity i*n vivo*. From an unbiased quantitative global phosphoproteomics analysis we identified several novel Orb6 targets related to membrane trafficking, including proteins implicated in signaling, lipid binding, and exocytosis. We confirmed Orb6 phosphorylation of exocyst proteins (Sec3, Sec5, and Sec8) and demonstrated the importance of Sec3 serine-201 phosphorylation in exocytosis.

### Role of Gef1 in mediating Orb6 function

Orb6 inactivation, using either temperature-sensitive *orb6-25* or analog-sensitive *orb6-as2*, results in several phenotypes, including (but not limited to): some ectopic localization of Cdc42-GTP to cell sides, actin disorganization, and increased cell width and/or rounding. Cdc42, a conserved Rho-family GTPase, is fundamental to multiple aspects of cell polarity, including actin organization and exocyst function^47^, and thus it has been proposed that ectopic Cdc42-GTP is a critical mediator of *orb6* phenotypes^25^. However, our results indicate that ectopic Cdc42-GTP localization is not the cause of increased cell width in *orb6* cells, because cell-width increases after Orb6 inactivation in wild-type (*gef1+*) cells are similar to those seen in *gef1∆* cells, which do not have ectopic Cdc42-GTP. Our results disagree with earlier work reporting smaller width increases in *gef1∆* compared to *gef1+* cells after Orb6 inactivation^25^. Currently we do not have an explanation for these different findings, which led us to investigate alternative mechanisms by which Orb6 could regulate polarized growth (see below).

We note that that the cell-width increase in *orb6-25* mutants at restrictive temperature is much greater than in Orb6-inhibited *orb6-as2* cells (Fig. 1C). The reason for this difference is unknown, but it could indicate kinase-independent functions of Orb6; it is also possible that Orb6-25 protein could be degraded at restrictive temperature, which might could affect the behavior or stability of Orb6-interacting proteins.

### Orb6 and exocytosis

Orb6-inhibited cells (both *gef1+* and *gef1∆*) are strongly impaired in exocytosis and cease interphase cell elongation, in spite of enriched Cdc42-GTP at cell tips. The exocytosis and elongation phenotypes are likely related, as illustrated by the disappearance of beta-glucan synthase Bgs4 from cell tips after Orb6 inhibition. Bgs4 disappearance is due to continued endocytosis in the absence of exocytosis, and because Bgs4 is required for cell wall synthesis at cell tips, its disappearance from cell tips can account, at least in part, for cessation of cell elongation.

In Orb6-inhibited cells, although exocytic trafficking of Bgs4 to cell tips is abolished during interphase, Bgs4 nevertheless accumulates at the cell midzone, on either side of the primary septum, during cytokinesis. This indicates that during cytokinesis, exocytosis proceeds independently of Orb6, to facilitate processes such as primary septum formation. Later, as cells progress to the next interphase, Orb6-dependent exocytosis is required for secretion of hydrolases to dissolve the primary septum and separate the two daughter cells. Our finding that Orb6 regulates exocyst (see below) is consistent with the fact that exocyst mutants display cell-separation defects rather than cytokinesis defects^34, 45, 46^. This in turn implies that targeting of secretion towards the midzone during cytokinesis (e.g. for primary septum formation) involves mechanisms other than exocyst^34^. Based on previous work showing redundancy of exocyst function and actin cable-based vesicle delivery in promoting efficient exocytosis^48–50^, we speculate that during cytokinesis, the abundance of actin cables contributing to the cytokinetic actomyosin ring^51, 52^ may bypass any requirement for exocyst.

The cell-separation defect of Orb6-inhibited cells is suppressed by deletion of *sts5*, which encodes an RNA-binding protein^23^. It has been proposed that Orb6 negatively regulates the recruitment of Sts5 into ribonucleoprotein granules and P-bodies, thereby controlling the translational repression of Sts5-associated mRNAs encoding proteins involved in polarized growth and cell separation^23^. Because our data suggests that cell-separation defects in Orb6-inhibited cells are due to defects in exocytosis, it is plausible that *sts5∆* causes an upregulation in exocytosis post-cytokinesis, promoting dissolution of the primary septum and thus cell separation. Potential links between Sts5 and exocytosis represent an interesting area for future exploration.

### Phosphoregulation of exocyst

Through quantitative global phosphoproteomics, phosphomapping by immunoprecipitation, and *in vitro* kinase assays we identified Orb6-dependent phosphorylation of exocyst proteins Sec3, Sec5, and Sec8. Analysis of *sec3-S201A* phenotypes suggests that the function of Sec3 serine-201 phosphorylation overlaps with that of Exo70. Thus far it remains unclear precisely what this function is. Our MS data suggest that exocyst remains largely intact upon Orb6 inhibition. Moreover, in *sec3-S201A exo70∆* cells, both Sec5 and Sec8 can localize to cell tips and the septation zone. In both budding and fission yeast, Sec3 and Exo70 interact with PIP2 on the plasma membrane, and this interaction is required (redundantly; either interaction is sufficient) for plasma-membrane localization of other exocyst proteins^45, 53–55^. Together with the fact that fission yeast *sec3∆ exo70∆* double mutants are lethal^45^, this suggests that *sec3-S201A* is a partial rather than a complete loss-of-function allele. We speculate that Sec3 serine-201 phosphorylation may influence exocyst function by modulating its conformational dynamics^56^ and/or interaction with small GTPases^57^. Exocyst is regulated by Rho- and Rab-family small GTPases^57^ as well as by protein kinases^28^, and this regulation can alter exocyst complex assembly and/or local activation of exocyst. Fission yeast Sec3 has an N-terminal cryptic PH domain and a C-terminal predicted coiled-coil domain^45^; however, serine-201 lies outside of these domains.

### Additional Orb6 targets

Orb6 inhibition *in vivo* leads to cessation of cell elongation, whereas *sec3-S201A exo70∆* cells, in spite of their cell-separation defect, can nevertheless grow in a polarized fashion (Fig. 6). We propose that the strong phenotypes seen after Orb6 inhibition are due to the combined changes in phosphorylation of multiple Orb6 targets, in multiple related pathways. In addition to Sec3 and Sec5, Orb6 targets identified here include protein kinases and proteins involved in membrane trafficking. Nearly all of the highest-ranking phosphosites (i.e. those with highest reproducibility and greatest fold-decrease after Orb6 inhibition) perfectly match the NDR/LATS consensus (Fig. 4, Suppl. Fig. 2, Suppl. Table 1), suggesting that they are direct substrates of Orb6.

In particular, in addition to exocyst, the proteins Sec72, Pep7, Ppk25 and Osh2 suggest a significant role for Orb6 in membrane trafficking. Sec72 is a homolog of budding yeast Sec7, a GEF for Arf1 ADP ribosylation factor, important for coat formation on nascent vesicles emanating from the Golgi and Trans-Golgi Network^58^. Budding yeast Pep7 (also called Vac1/Vps19) integrates Rab GTPase and PI3-kinase signaling and promotes Golgi-to-endosome transport^**59**^. The kinase Ppk25 does not have extensive regulatory domains, but its kinase domain is most similar to that of the budding yeast paralogs Kin1 and Kin2, which function in late stages of exocytosis through phosphorylation of t-SNARE Sec9^60^. Budding yeast Osh2, a member of a conserved family of oxysterol-binding protein-related proteins^61^, facilitates endocytosis at plasma membrane regions that contact cortical endoplasmic reticulum^62^. Collectively, the functions of these proteins in budding yeast suggest that in fission yeast, Orb6 coordinates multiple processes related to membrane trafficking. In this context, we note that while we identified several phosphosites in both Gef1 and Sts5, none of these showed significantly decreased phosphorylation (i.e. two-fold or greater) after Orb6 inhibition (Suppl. Table 2). It is thus unclear whether Gef1 or Sts5 are bona fide Orb6 substrates *in vivo*. However, it should be recognized that our global phosphoproteomics analysis is unlikely to be exhaustive.

The notion that Orb6 may have multiple roles in regulating membrane trafficking and cell polarity is consistent with analysis of NDR1/2 kinase function in mammalian cells. NDR1 negatively regulates dendrite length and branching via phosphorylation of targets including AAK1 (AP-2 associated kinase) and Rabin8 (GEF for GTPase Rab8), which regulate vesicle-mediated trafficking^20^. Independently, NDR2 was shown to phosphorylate Rabin8 to promote Rabin8-Sec15 interaction, activating Rab8 during ciliogenesis^63^. NDR2 also regulates β1-integrin exocytosis, thus controlling integrin activation for neurite growth and guidance^64^. In addition, in budding yeast, Cbk1 has been proposed to regulate more than one step in the secretory pathway^22^, although some aspects of this may be controversial^65^. Collectively, our results suggest that regulation of exocytosis/secretion by NDR kinases may be a generally conserved design principle, although specific targets of NDR kinases may differ in different systems.

### Physiological function of switching off Orb6 activity

Orb6 is the most downstream kinase of the fission yeast Morphogenesis Network (MOR), a signaling pathway similar in design to the fission yeast Septation Initiation Network (SIN)^66^, budding yeast MEN and RAM^21^, and metazoan Hippo^17^ pathways. In addition to Orb6, MOR proteins include Sog2, Mob2, Mor2 and Pmo25 as well as the germinal-centre kinase Nak1; all of these are conserved, and all are important for the regulation of cell polarity and cell separation^67^. MOR signaling decreases during mitosis, when cells switch polarity from tip growth to the midzone for cell division, and it increases again after completion of cytokinesis, coincident with cell separation^67, 68^. Attenuation of MOR signaling in mitosis depends on SIN, and there is compelling evidence that mutual antagonism between MOR and SIN is critical for coordinating changes in cell polarity and cytoskeletal reorganization through the cell cycle^68^. At the same time, however, the actual targets of MOR that execute these changes have generally remained obscure. Based on the results presented here, we propose a simple model in which Orb6-regulation of exocytosis is a key output of MOR signalling. According to this view, during interphase, Orb6 would function to promote polarized exocytosis at cell tips. During mitosis, decreased Orb6 activity would shut down exocytosis at tips, allowing Orb6-independent redistribution of secretion towards the cell midzone. After cytokinesis (effectively, the next interphase), increased Orb6 activity would specifically promote exocytosis of hydrolases into the septation zone, dissolving the primary septum for daughter-cell separation and allowing resumption of growth at cell ends. While further functional characterization of Orb6 targets awaits elucidation, we speculate that by coordinating multiple kinase signaling and vesicle-mediated trafficking pathways, Orb6 may function as an interphase-specific master regulator of membrane trafficking and exocytosis, thus helping to drive transitions in cellular organization between interphase and mitosis.

## MATERIALS AND METHODS

### Yeast culture

Standard fission yeast methods were used throughout^69, 70^. Growth medium was either YE5S rich medium (using Bacto yeast extract; Becton Dickinson) or Edinburgh Minimal Medium. Supplements such as adenine, leucine, and uracil were used at 175 mg/L. Solid media used 2 % Bacto agar (Becton Dickinson). Normal arginine and lysine supplements were used at 40 mg/L and 30 mg/L, respectively. “Heavy” L-arginine-HCl (^13^C_6_-labelled; Sigma; No. 643440) and L-lysine-HCl (^13^C_6_-labelled, Cambridge Isotope laboratories) supplements were used at the same concentration as their light counterparts. SILAC cell cultures were grown as described previously^39, 40^. To inhibit Orb6 kinase activity *in vivo*, 3-BrB-PP1 (Toronto Research Chemicals; A602985), 3-MB-PP1 (Toronto Research Chemicals; A602960), 1-NM-PP1 (Toronto Research Chemicals; A603003) dissolved in methanol were each used at a final concentration of 30 µM. For imaging experiments involving hydroxyurea, cells were grown in YE5S media containing 12 mM hydroxyurea (prepared fresh; Sigma; H8627), starting at 1.4 hr before the beginning of imaging. For actin depolymerization, Latrunculin A (Alpha laboratories; 129-04361) was added to cells at a final concentration of 50 µM in growth medium (made from a 25 mM stock in DMSO), 10 min before addition of 3-BrB-PP1.

### Plasmid and yeast strain construction

Mating for genetic crosses^71^ was performed on SPA5S plates with supplements at 45 mg/L. Crosses were performed using tetrad dissection or random spore analysis. Tagging and deletion of genes were performed using PCR-based methods^72^. All strains used in this study are listed in Supplementary Table 3.

### Construction of orb6-as2 strain

Two DNA fragments encoding N- and C-terminal regions of Orb6 were amplified from fission yeast genomic DNA. Both DNA sequences contain 25 nucleotide overlapping sequence and a single point mutation (*orb6-as2*) corresponding to an M170A mutation in the amino-acid sequence. The two fragments were assembled with plasmid pREP41X using Gibson assembly kit (NEB; E2611S), to make plasmid pKS1439. An NdeI-BamHI restriction fragment of pKS1439 containing the mutated coding sequence was transformed into an *orb6-25* strain. Selection was performed at 37ºC, which is non-permissive for the growth of *orb6-25* cells. Positive clones were sequenced to ensure correct replacement of the *orb6-25* allele by *orb6-as2*.

### Microscopy sample preparation and imaging

Exponentially growing cells cultured at 25°C were used in all imaging experiments, unless otherwise stated. Fluorescence live-cell imaging was performed either in coverslip dishes (MatTek; P35G-0.170-14-C.s) or 4-chamber glass bottom micro-slides (Ibidi; 80427). Imaging dishes/slides were coated with 1 mg/mL soybean lectin (Sigma; L1395) for 10 min and washed with appropriate medium to remove excess lectin. Log-phase culture was added to dishes/slides and allowed to attach to the coverslip bottom for 15 min. The dishes/slides were washed extensively with media using aspiration with at least 3 full exchanges of media (approximately 1 ml each). Finally, 500 µL of medium was added to the dish/slide before imaging. Nearly all live-cell fluorescence imaging was done using a custom spinning-disk confocal microscope unit composed of Nikon TE2000 microscope base, attached to a modified Yokogawa CSU-10 unit (Visitech) and an iXon+ Du888 EMCCD camera (Andor), 100x/1.45 NA Plan Apo objective (Nikon), Optospin IV filter wheel (Cairn Research), MS-2000 automated stage with CRISP autofocus (ASI), and thermo-regulated chamber maintained at 25°C (OKOlab). Metamorph software (Molecular Devices) was used to control the spinning-disc confocal microscope. FRAP was performed on a Zeiss Airyscan microscope (LSM880, AxioObserver, alpha Plan-ApoChromat 100x/1.46 Oil DIC M27 Elyra). GFP signal was detected using 488 nm line of an Argon laser and a 495-550 nm bandpass emission filter (number of iteration=5, acquisition time interval 30s).

Calcofluor staining was performed according to standard protocol^73^. Briefly, 900 µL of log-phase culture was added to 100 µL 30 % formaldehyde solution (Sigma; F1268) and kept on ice for 10 min. Cells were pelleted and washed three times using ice-cold PBS buffer. The final pellet was resuspended in 50 µL PBS. 1.5µL of cell suspension was mixed with 1.5 µL of 10 µg/mL Calcofluor solution (Fluorescence Brightener 28; Sigma; F3543). Calcofluor-stained cells were imaged using a Zeiss Axiolmage microscope with 100X/1.46NA alpha-Plan Apochromat objective, CoolEd Pe300 lightsources and Hamamatsu Flash 4 camera. Micromanager acquisition software was used to control the Zeiss microscope.

### Microscopy image analysis

ImageJ (Fiji, NIH) was used to process all acquired raw images. All images and movies shown are maximum projections of eleven Z-sections with 0.7 μm step-size. ImageJ StackReg plugin was used for rigid body registrations. Kymographs was generated using Reslice and KymoResliceWide plugin in ImageJ. Cell-width and cell-elongation measurements were performed manually using line tool in ImageJ. Image formatting and assembly were performed using Photoshop (Adobe) and Illustrator CS3 (Adobe). Movies were edited using ImageJ and QuickTime (Apple). Graphs were created using Graphpad Prism software and Microsoft Excel.

### Acid phosphatase assay

Acid phosphatase assay was performed as described previously^34^. Cells were grown to mid-log phase in YE5S at 25ºC. Cells were washed twice in pre-warmed YE5S medium, and all strains were adjusted to OD_595_=0.25. The cells in fresh media were cultured in a 25ºC shaking water bath. At indicated time points, cell density (OD_595_) was measured, and 1 mL culture was centrifuged to pellet the cells. 300 µL cell-free culture medium was incubated with 400 µL of phosphatase substrate solution (3 tablets of phosphatase substrate (Sigma; S0942) dissolved in 20 mL of 0.1 M sodium acetate pH 4.0, pre-warmed to 32ºC). The reaction was stopped by addition of 400 µL of 1 M NaOH. OD_405_ was measured against blank consisting of fresh YE5S medium without cells. For each time point, ACP activity (OD_405_)was normalized to cell density (OD_595_). In the experiments in Fig. 3B, four independent biological replicates were performed for wild-type and *orb6-as2*, and two independent biological replicates for *orb6-as2 gef1∆.* In the experiments in Fig. 6D, three independent replicates were performed for *exo70∆* and *exo70∆ sec3-S201A*, and two independent biological replicates for *exo70∆ sec5-S50A*.

### Sample preparation for global phosphoproteomics analysis

SILAC cultures supplemented with heavy arginine and lysine were grown for at least 8 generations to ensure complete isotopic labelling. Light and heavy SILAC cultures were grown to OD_595_ 0.75 and then treated with either methanol or 30 µM 3-BrB-PP1 for 3 hr. In two replicate experiments, heavy cultures were treated with 3-BrB-PP1 and light cultures were treated with methanol, and in a third replicate, heavy cultures were treated with methanol and light cultures were treated with 3-BrB-PP1. Cells were harvested by centrifugation and washed once with STOP buffer (10 mM EDTA, 50 mM NaF, 150 mM NaCl, 1 mM NaN_3_). The pelleted cells were resuspended in milliQ H_2_O at 400 mg/mL concentration and flash-frozen in liquid nitrogen. Mechanical lysis of frozen cells (cryogrinding) was performed under liquid nitrogen in a SPEX SamplePrep LLC 6870 Freezer/mill^®^ (sample pre-cool for 2 min, 10 rounds of “run” and “cool” cycle each for 2 min, beat rate=10). ~800 mg of light and heavy cell powders were individually solubilized in denaturation buffer (6 M urea, 2 M thiourea, 10 mM Tris-HCl pH 8.0) for 1 hr at room temperature and centrifuged at 4,500 x *g* for 15 min to obtain a clear supernatant. Protein concentration from both samples was measured using Bradford assay.

For protein abundance measurement of the SILAC samples, light and heavy cell lysates (5 mg each) were mixed, and 50 µg of the mixed protein lysate was separated by SDS-PAGE on Bolt^®^ 4-12 % Bis-Tris Gel (Thermo Fisher Scientific; NW04120BOX). The protein gel was stained by Coomassie brilliant blue (Sigma; B0149) and the entire gel lane was excised into 10 bands and processed for protein abundance analysis (see “In-gel digestion for mass spectrometry”, below).

For SILAC phosphopeptide quantification, the remaining 9.95 mg of the mixed protein lysate was reduced in 1 mM DTT for 1 hr at room temperature. The reduced lysate was further alkylated in 5.5 mM iodoacetamide (Sigma; I1149) for 1 hr at room temperature in the dark. In-solution protein digestions were performed using Lysyl Endopeptidase (LysC) and trypsin in two stages. First, the lysate was digested by 90 µg of endopeptidase LysC (Waco Chemicals; 129-02541) for 3 hr at room temperature. The lysate was then diluted 5 times to final concentration of 1.6 M urea/ 0.4 M thiourea. 90 µg of trypsin (Thermo-scientific; 90057) was added to the lysate and incubated for 24 hr at room temperature. Digestion was terminated by acidification to final concentration of 0.4 % trifluoroacetic acid (TFA) (Sigma; 6508).

### Peptide fractionation using strong cation exchange (SCX) chromatography

SCX fractionation was performed as in^74^ with some modifications. Doubly-digested mixed lysate was fractionated using a Resource S SCX column (1 mL, GE Healthcare) in a ÄKTA protein purification system (GE Healthcare). Briefly, the mixed lysate was loaded onto the SCX column equilibrated in Solvent A (5 mM potassium dihydrogen phosphate, 30 % acetonitrile (ACN) (Fisher Chemicals, A955-212), pH 2.7 with TFA) at a flow rate of 1 mL/min. Flow-through during loading was kept for subsequent phosphopeptide enrichment. Peptides bound to the column were eluted as 2 mL fractions with a 0-50 % linear gradient of Solvent B (350 mM potassium chloride, 5 mM potassium dihydrogen phosphate, 30 % ACN, pH 2.7 using TFA) at a flow rate of 1 mL/ min, over a period of 30 min. Dilute fractions, based on estimation from the chromatogram, were pooled together for subsequent phosphopeptide enrichment.

### Phosphopeptide enrichment using titanium dioxide (TiO2) beads

TiO_2_ phosphopeptide enrichment was performed as in^75^ with some modification. Each SCX fraction, including the flow-through and pooled fractions, was processed individually. Flow-through fraction was incubated with 10 mg TiO_2_ beads slurry (Titansphere; GL Sciences; 5020-75010) in 30 mg/mL 2,5 dihydrobenzoic acid (Sigma; 149357) and 80 % ACN, whereas other fractions were incubated with 5 mg TiO_2_ beads slurry and incubated at room temperature for 30 min. TiO_2_ beads were concentrated by 1 min centrifugation at 13,000 rpm and supernatant was removed. TiO_2_ beads were then washed with washing solution I (30 % ACN, 3 % TFA) and pelleted. TiO_2_ beads were then re-suspended with washing solution II (80 %, ACN, 0.1 % TFA) and transferred to C8-stage tip (~1 mm^2^ Empore C8 in 200 µL pipette tip). Washing solution II was removed from the stage-tip by centrifugation at 4,000rpm for 2-3 min. Phosphopeptides bound to the TiO_2_ beads were eluted three times by passing through elution solution (40 % ammonium hydroxide (Sigma; 338818), 60 % ACN). The eluates were loaded onto C18-stage tip (~1mm^2^ Empore C18 in 200 µL pipette tip) pre-washed with 0.1 % TFA. The stage tips were washed with 50 µL 0.1

% TFA and stored at −20ºC until further use. Because the flow-through fraction contains the most abundant phosphopeptide fraction, two additional rounds of phosphopeptide enrichment were performed on the flow-through fraction with fresh TiO_2_ beads (10 mg).

### In-gel digestion for mass spectrometry

Coomassie-stained gel slices were destained and digested in-gel as described^76^. Briefly, the bands were destained by 3-4 rounds of incubation in 25 mM ammonium bicarbonate in 50 % ACN until the last trace of Coomassie blue was removed. Gel slices were reduced in 10 mM DTT for 30 min at 37ºC, alkylated in 55 mM iodoacetamide for 30 min at room temperature in the dark, and digested overnight at 37ºC with 12.5 ng/µL trypsin (Thermo-scientific; 90057). Digestion medium was then acidified by TFA and the eluted peptides were desalted using C18-stage tip as described^77^.

### LC-MS/MS

Peptides were eluted in 40 μL of 80 % acetonitrile in 0.1 % TFA and then concentrated down to 1μl by vacuum centrifugation (Concentrator 5301, Eppendorf, UK). Samples were then diluted to 5 µl with 0.1 % TFA and prepared for LC-MS/MS analysis. LC-MS-analyses were performed on a Q Exactive mass spectrometer (Thermo Fisher Scientific, UK) and on an Orbitrap Fusion™ Lumos™ Tribrid™ Mass Spectrometer (Thermo Fisher Scientific, UK). Both were coupled on-line to Ultimate 3000 RSLCnano Systems (Dionex, Thermo Fisher Scientific, UK). For samples that were analysed by Q Exactive, peptides were separated by an analytical column with a self-assembled particle frit (Ishihama *et al* 2002) and C18 material (ReproSil-Pur C18-AQ 3 μm; Dr. Maisch, GmbH, Germany) that was packed into a spray emitter (75 μm ID, 8 μm opening, 300 mm length; New Objective) using an air-pressure pump (Proxeon Biosystems, USA). The column was maintained at stable temperature (40°C) with the appropriate column oven (Sonnation, Germany). Peptides that were analysed by Fusion Lumos were separated on a 50cm EASY-Spray column (Thermo Scientific, UK) assembled in an EASY-Spray source (Thermo Scientific, UK) and operated at 50°C. In both cases, mobile phase A consisted of 0.1 % formic acid in water while mobile phase B consisted of 80 % acetonitrile and 0.1 % formic acid. Peptides were loaded at a flow rate of 0.5 μl min^−1^ (for samples that analysed by the Q Exactive) and 0.3 μl min^−1^ (for samples on Fusion Lumos) and eluted at a flow rate was 0.2 μl min^−1^according to the following gradient: 2 to 40 % buffer B in 120 min, then to 95 % in 11 min.

For Q Exactive, FTMS spectra were recorded at 70,000 resolution (scan range 300-1700 m/z) and the ten most intense peaks with charge ≥ 2 of the MS scan were selected with an isolation window of 2.0 Thomson for MS2 (filling 1.0E6 ions for MS scan, 5.0E4 ions for MS2, maximum fill time 60ms, dynamic exclusion for 50 s). For Orbitrap Fusion™ Lumos™, survey scans were performed at 120,000 resolution (scan range 400-1900 m/z) with an ion target of 4.0e5. MS2 was performed in the ion trap with ion target of 1.0e4 and HCD fragmentation with normalized collision energy of 27^78^. The isolation window in the quadrupole was 1.4. Only ions with charge between 2 and 7 were selected for MS2.

The MaxQuant software platform^79^ version 1.5.2.8 was used to process raw files, and search was conducted against *Schizosaccharomyces pombe* complete/reference proteome set of Pombe database (released in August, 2013), using the Andromeda search engine^80^. The first search peptide tolerance was set to 20ppm while the main search peptide tolerance was set to 4.5 pm. Isotope mass tolerance was 2 ppm and maximum charge to 7. Maximum of two missed cleavages were allowed. Carbamidomethylation of cysteine was set as fixed modification. Oxidation of methionine, acetylation of the N-terminal and phosphorylation of serine, threonine and tyrosine were selected as variable modifications. When SILAC labelled samples were analysed, the multiplicity was set to 2 and the appropriate labels were selected. Peptide and protein identifications were filtered to 1 % FDR.

### Immunoprecipitation of Sec5-3mCherry

Sec5-3mCherry cells in *SILAC* background were grown in two separate cultures supplemented with light and heavy isotopes for 8 generations. Light and heavy cultures were treated with methanol and 30 µM 3-BrB-PP1 respectively, for 45 min. The cells were washed with STOP buffer and resuspended in lysis buffer (50 mM Tris-HCl pH 7.5, 50 mM NaF, 150 mM NaCl, 20 mM Na-β-glycerophosphate, 0.2 % Triton-X100, 1 mM Na_3_VO_4_, 1 mM EDTA, 10 μg/mL each of ‘CLAAPE’ protease inhibitors, 2 mM AEBSF, 2 mM benzamidine, 50 nM Calyculin A, 50nM Okadaic acid and 2 mM PMSF) at a concentration of 1 g/mL. The cell suspension was drop-frozen in liquid nitrogen and mechanically lysed using Freezer/Mill^®^. Light and heavy cell powders were resuspended in lysis buffer and cell extracts were cleared by 2 × 15 min centrifugation at 4,000 x *g* in a Beckman Avanti J-26 centrifuge using a JLA 8.1000 rotor. Protein content in each sample was measured and samples are mixed at 1:1 ratio. The mixed cell extract was incubated with Protein G Dynabeads (Thermo Fisher Scientific), covalently crosslinked to homemade affinity-purified rabbit anti-tdimer2 using dimethyl pimelimidate, for 1 hr. Dynabeads were then washed five times with lysis buffer. Proteins were eluted from Dynabeads by 10 min incubation at 65ºC in Laemmli sample buffer. Eluted proteins were separated by SDS-PAGE on a Bolt^®^ 4-12 % Bis-Tris Gel. The gel was stained by Coomassie Brilliant Blue, and gel bands were excised and processed according to the in-gel digestion protocol (see above).

### Purification of recombinant GST-Sec3 and GST-Sec5 proteins

Sec3 and Sec5 sequences amplified from fission yeast genomic DNA were cloned into pGEX4T-1 vector and expressed as GST fusion proteins in *E.coli* strain BL21(DE3RIL). Expression was induced in 500 mL bacterial culture at OD_595_=0.8 by addition of 0.2 mM IPTG, followed by growth for additional 24 hr at 18ºC. Cells were harvested by centrifugation and resuspended in lysis buffer (50 mM Tris-HCl pH 8.0, 150 mM NaCl, 5 % glycerol, 5 mM EDTA, 3 mM DTT, 0.1 % Triton-X100, 200 µg/mL lysozyme, benzonase (10 units/mL), 10 μg/mL each of ‘CLAAPE’ protease inhibitors). The cells were then lysed by sonication. Cell extracts were clarified by centrifugation at 15,000 rpm at 4ºC for 30 min. Cleared lysates were then incubated with glutathione-agarose (Sigma; G4510). Glutathione-agarose with bound GST proteins was then placed in a column and washed with 20 bead-volumes of wash buffer (50 mM Tris-HCl pH 8.0, 300 mM NaCl, 100 mM KCl, 10 % glycerol, 5 mM EDTA, 3 mM DTT, 0.1 % Triton-X100). GST fusion proteins were then eluted with elution buffer (50 mM Tris-HCl pH 8.0, 300 mM NaCl, 10 % glycerol, 2 mM EDTA, 3 mM DTT, 20 mM glutathione) in 1 mL fractions. Protein-rich fractions were pooled together and concentrated using Amicon Ultra-4 centrifugal filter unit with Ultracel^®^-30 membrane (Millipore; UFC803024).

### Mob2 In vitro kinase using GST-Sec3 and GST-Sec5

Mob2-GFP cells in *sts5∆* and *sts5∆ orb6∆* backgrounds were grown in YE5S medium to mid-log-phase, harvested and washed with STOP buffer. 1×10^7^ Cells were pelleted into 1.5 mL screw-cap microfuge tubes and snap frozen in liquid nitrogen. 200 µL of precooled lysis buffer (50 mM Tris-HCl pH 7.5, 50 mM NaF, 150 mM NaCl, 20 mM Na-β-glycerophosphate, 0.2 % Triton-X100, 1 mM Na_3_VO_4_, 1 mM EDTA, 10 μg/mL each of ‘CLAAPE’ protease inhibitors, 2 mM AEBSF, 2 mM benzamidine, 50 nM calyculin A, 50 nM okadaic acid and 2 mM PMSF) and ~500 µL of 0.5 mm zirconia beads were added to the microfuge tubes and disrupted by bead beating in Ribolyser (Hybaid) at 4ºC using two cycles of 35 s at maximum setting (6.5). 350 µL of lysis buffer was added into the tube and mixed thoroughly. Cell lysates were collected by centrifugation through a hole pierced at the bottom of the microfuge tube. Cell lysates were cleared by centrifugation at 13,000 rpm for 6 min at 4ºC. 550 µL of cell lysates were incubated with 10 µL of Protein G Dynabeads, coupled to homemade affinity-purified Sheep anti-GFP, for 1 hr at 4ºC. Dynabeads were then washed 3 times using lysis buffer and 3 times using kinase buffer (50 mM Tris-HCl pH 7.5, 100 mM NaCl, 10 mM MgCl_2_, 1 mM MnCl_2_, 10 mM ATP, 20 mM Na-β-glycerophosphate, 1 mM Na_3_VO_4_, 10 μg/mL each of ‘CLAAPE’ protease inhibitors). Dynabeads were then incubated with 5 µL of recombinant GST-Sec3 and GST-Sec5 in 20 µL kinase buffer at 30ºC for 30 min. The kinase reaction was stopped and recovered from Dynabeads by incubation in Laemmli sample buffer at 65ºC for 10 min. Kinase reactions and controls were separated on Bolt^®^ 4-12 % Bis-Tris gel and stained with Coomassie. Bands corresponding to GST-Sec3 and GST-Sec5 were excised and processed according to in-gel digestion protocol.

### Phosphosite ranking and GO analysis

Phosphosites with a two-fold or greater mean decrease in phosphorylation after Orb6 inhibition are presented in Suppl. Table 1, which shows three categories of stringency. Phosphosites in the “lowest stringency” category were quantified in at least one of three replicate experiments, while phosphosites in the “medium stringency” category were quantified in at least two out of three replicate experiments, and phosphosites in the “highest stringency” category were quantified in all three replicate experiments. Within each category, phosphosites were ranked by mean fold-decrease in phosphorylation after Orb6 inhibition (largest negative values correspond to greatest fold-decrease).

Alignment of sequences containing the NDR/LATS consensus was performed using MegAlignPro version 15 (DNASTAR). GO analysis was carried out using Generic Gene Ontology (GO) Term Finder^81^. Fold-enrichment for each GO term identified from our input cluster of genes is defined as the ratio of (frequency of the GO term within the cluster) to (frequency of the GO term within the genome).

### Statistical tests

Statistical analyses in Figs. 1C and 1E were determined by two-tailed unpaired Mann–Whitney test, using GraphPad Prism software (see Microscopy image analysis, above). Statistical significance in Fig. 4C was determined by Generic Gene Ontology (GO) Term Finder (see Phosphosite ranking and GO analysis, above).

## ACKNOWLEDGEMENTS

We thank Mohan Balasubramanian, Dannel McCollum, Taro Nakamura, Paul Nurse and Fulvia Verde, Pilar Perez, Juan Carlos Ribas and Hilary Snaith for strains, Iain Hagan and Avinash Patel for advice and protocols on phosphoproteomics, Harish Thakur and Su Ling Leong for advice on protein purification, and members of our laboratories for discussion. This work was supported by the BBRSC [BB/K021699/1 to K.E.S. and A.B.G.] and the Wellcome Trust [094517 to K.E.S.]. The Wellcome Centre for Cell Biology is supported by core funding from the Wellcome Trust [203149].

## SUPPLEMENTARY MOVIE LEGENDS

**Supplementary Movie 1. Orb6 inhibition leads to increased cell width, cessation of cell elongation and cell-separation defects in both wild-type and gef1∆ cells** Cdc42-GTP reporter CRIB-3mCitrine and F-actin reporter Lifeact-mCherry in *orb6-as2* and *gef1∆ orb6-as2* cells before and after 3-BrB-PP1 addition (indicated by “+3-BrB-PP1”). Note ectopic CRIB patches in *orb6-as2* cells but not *gef1∆ orb6-as2* cells after 3-BrB-PP1 addition. Time interval: 5 min. Total elapsed time: 415 min. Time compression at 15 frames per second playback: 4200x.

**Supplementary Movie 2. Cessation of interphase cell elongation after Orb6 inhibition** Cdc42-GTP reporter CRIB-3mCitrine in wild-type, *orb6-as2* and *gef1∆ orb6-as2* cells after 3-BrB-PP1 addition. 3-BrB-PP1 was added 1 hr before start of imaging. Note (faint) ectopic CRIB patches in *orb6-as2* cells but not *gef1∆ orb6-as2* cells after 3-BrB-PP1 addition. Movies correspond to cells shown in Fig. 1D. Time interval: 4 min. Total elapsed time: 96 min. Time compression at 15 frames per second playback: 3360x.

**Supplementary Movie 3. Loss of Bgs4 from cell tips after Orb6 inhibition.** mCherry-Bgs4 in exponentially-growing and hydroxyurea-arrested *orb6-as2* and *gef1∆ orb6-as2* cells before and after 3-BrB addition. Hydroxyurea was added 1.4 hr before start of imaging. Time interval: 4 min. Total elapsed time: 124 min. Time compression at 15 frames per second playback: 3360x.

**Supplementary Movie 4. Loss of Bgs4 from cell tips after Orb6 inhibition requires the actin cytoskeleton.** mCherry-Bgs4 in latrunculin A (LatA)-treated *orb6-as2* and *gef1∆ orb6-as2* cells before and after 3-BrB addition. LatA was added 10 min before start of imaging. Movie of *orb6-as2* cells contains a slight focus adjust. Time interval: 4 min. Total elapsed time: 140 min. Time compression at 15 frames per second play back: 3360x.

**Supplementary Figure 1.**
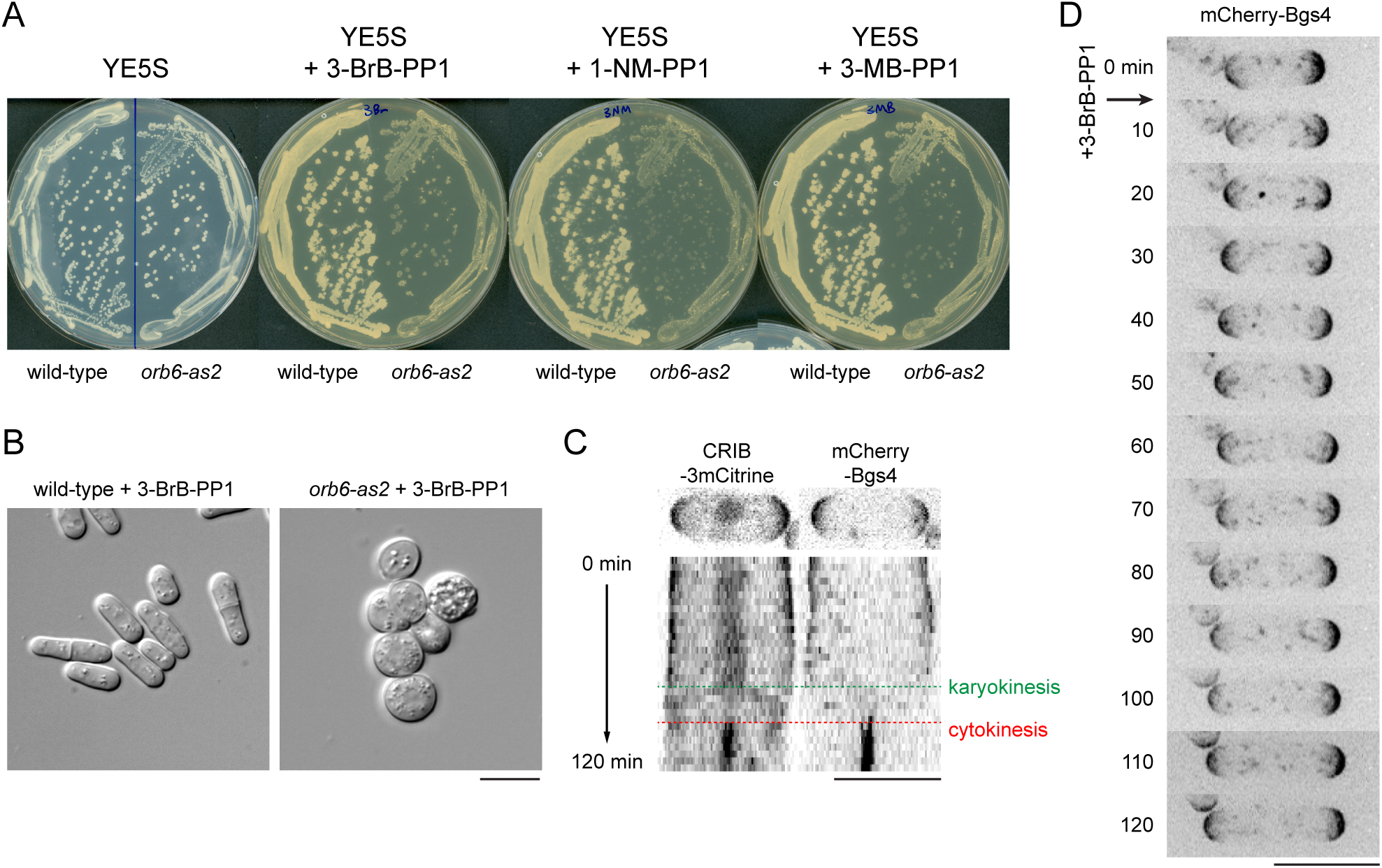
Additional characterization of *orb6-as2* and effects of 3-BrB-PP1. **(A)** Wild-type and *orb6-as2* colonies after replica-plating to the different indicated nucleotide-competitive analogs (each analog at 30 µM final concentration). All analogs inhibited growth of *orb6-as2* cells. Plates were incubated at 32ºC for 2 days. **(B)** DIC images of wild-type and *orb6-as2* cells taken from 3-BrB-PP1 plate. Note that Orb6-inhibited cells are rounder on solid media than in liquid medium (compare with Figure 1). **(C)** Temporal correlation of CRIB-3mCitrine and mCherry-Bgs4 localization to cell tips during interphase and to the division site during cell division. **(D)** Addition of 3-BrB-PP1 (30 µM) to wild-type cells does not affect mCherry-Bgs4 localization or interphase cell elongation. Bars, 10µm.

**Supplementary Figure 2.**
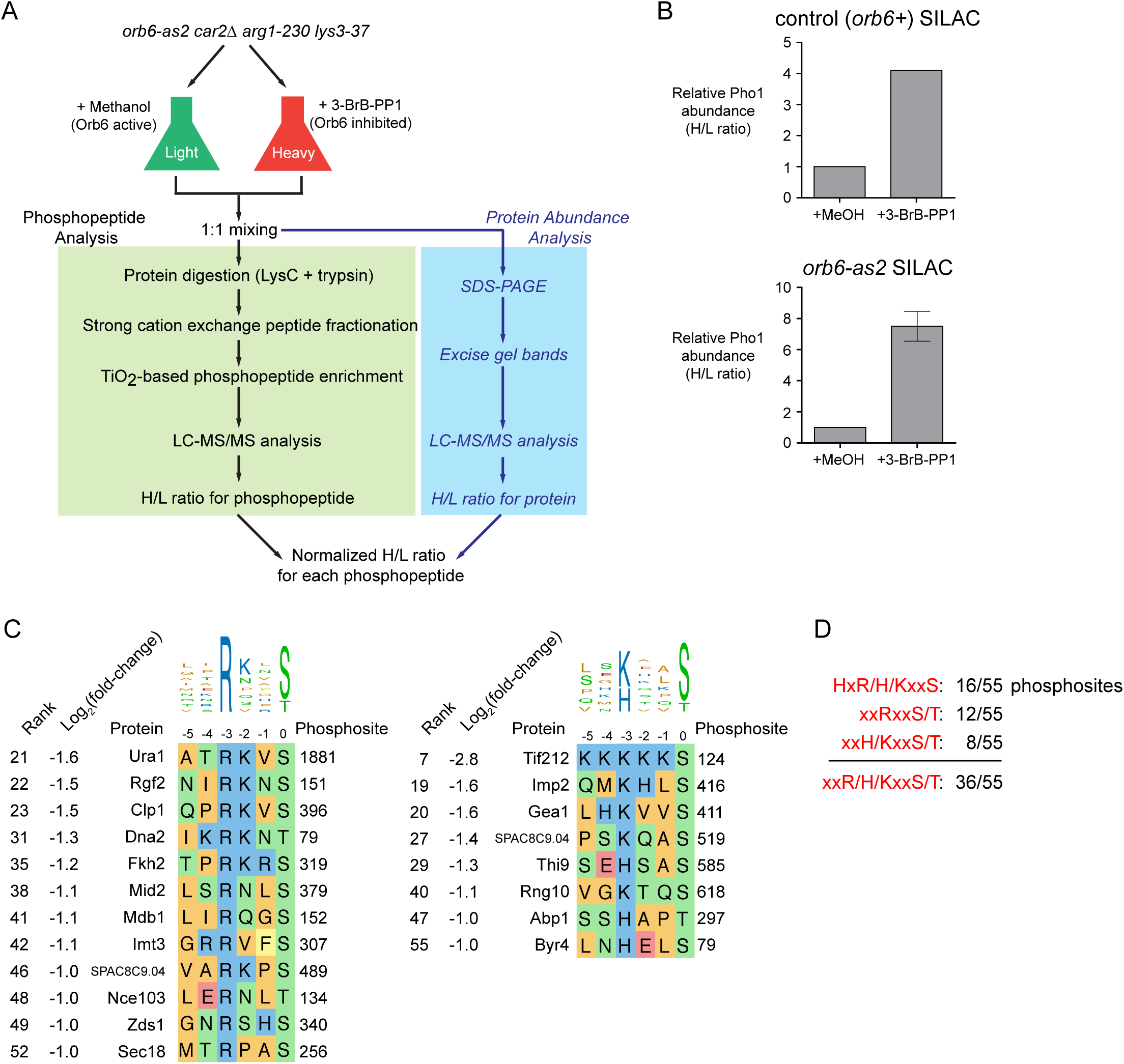
Quantitative global phosphoproteomics analysis after Orb6 inhibition. **(A)** Workflow for SILAC quantitative global phosphoproteomics analysis. Because relative abundance of individual proteins may change as a result of Orb6 inhibition, abundance of individual proteins before and after Orb6 inhibition is used to normalize H/L ratios of the relevant phosphopeptides. **(B)** Relative abundance of acid phosphatase Pho1 after 3-BrB-PP1 addition to control wild-type (*orb6+*) cells and *orb6-as2* cells. Error bar indicates SD from three biological replicates. Relative abundances of other acid phosphatases SPBC4.06 and Pho4 were not signficantly changed after 3-BrB-PP1 addition. **(C)** Alignment of selected highest-confidence Orb6-dependent phosphosites containing basic residues R or H/K at −3 position. All phosphosites shown were quantified in all three replicate global phosphoproteomics experiments and showed a two-fold or greater mean decreased phosphorylation after Orb6 inhibition. Phosphosites are ranked by fold-decrease in phosphorylation (see Fig. 4B and Suppl. Table 1). **(D)** Consensus summary of features of highest-confidence Orb6-dependent phospho-sites from (C), together with sites with perfect match to NDR/LATS consensus HxR/H/KxxS/T (Fig. 4B). 36 out of 55 sites include a basic residue at −3 position.

**Supplementary Figure 3.**
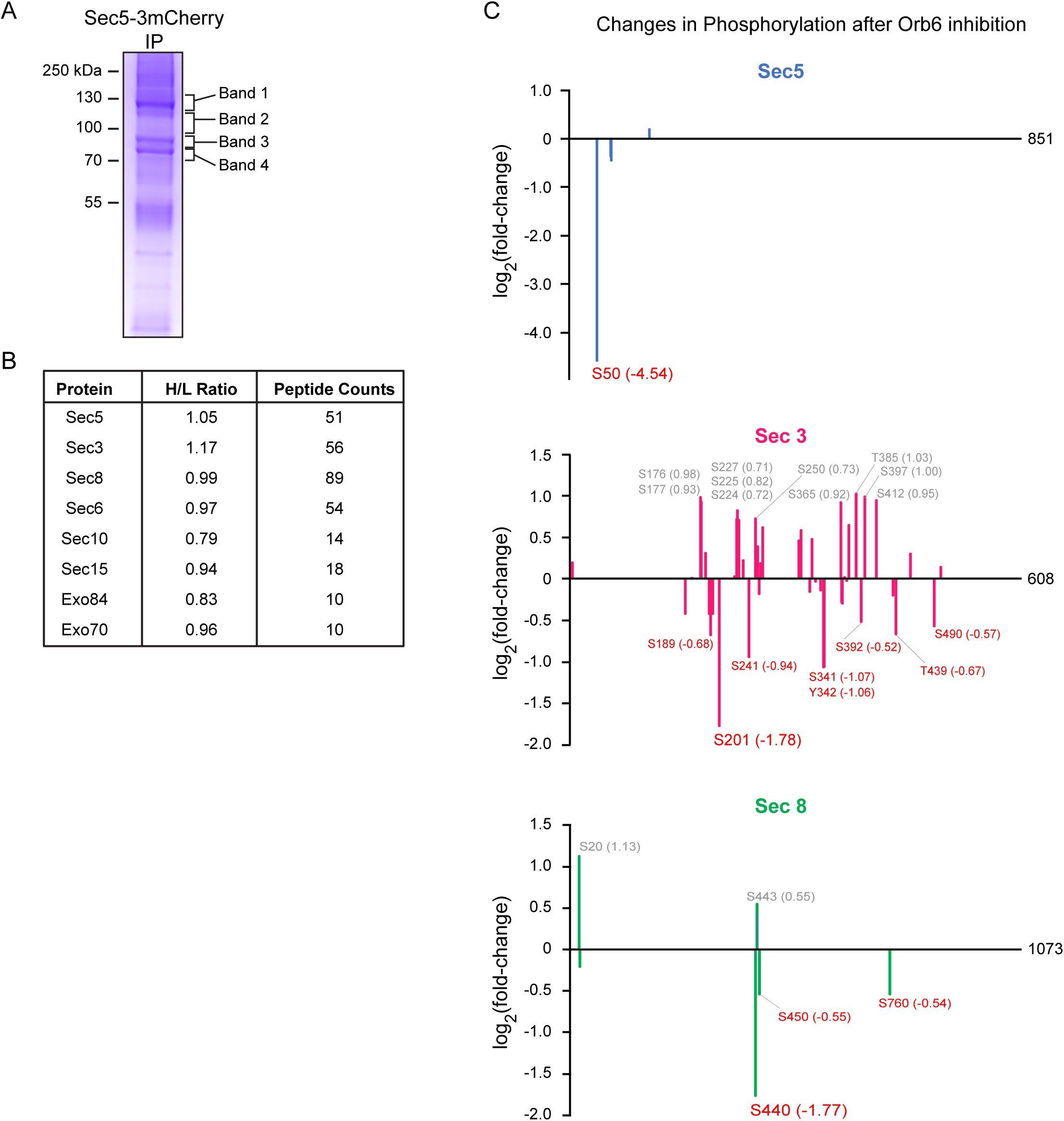
Additional confirmation of changes in phosphorylation of exocyst proteins after Orb6 inhibition. **(A)**SDS-PAGE showing immunoprecipitated Sec5-3mCherry and coimmunoprecipitated proteins, from a mixture of SILAC “heavy”-labelled Orb6-inhibited *orb6-as2* cells and “light”-labelled non-inhibited *orb6-as2* cells. Heavy-labelled cells were treated with 3-BrB-PP1, while light-labelled cells were treated with methanol. Indicated bands were excised and processed for tandem mass spectrometry. **(B)** Relative abundances of exocyst proteins present in SILAC Sec5-3mCherry immunoprecipitate from (A) before and after Orb6 inhibition, based on SILAC heavy:light ratio (H/L ratio). All ratios are close to 1, suggesting that Orb6 inhibition does not significantly alter exocyst integrity. **(C)** Fold-changes in phosphorylation of individual phosphosites in Sec5, Sec3, and Sec8 after Orb6 inhibition. X-axes show position of phosphosites in the respective proteins (Sec5 is 851 amino acids, Sec3 is 608 amino acids, and Sec8 is 1073 amino acids). Y-axes show log_2_(fold-change) in phosphorylation of relevant phosphosites after Orb6 inhibition. Among exocyst proteins, only Sec5, Sec3 and Sec8 showed significant changes in phosphorylation after Orb6 inhibition.

**Supplementary Figure 4.**
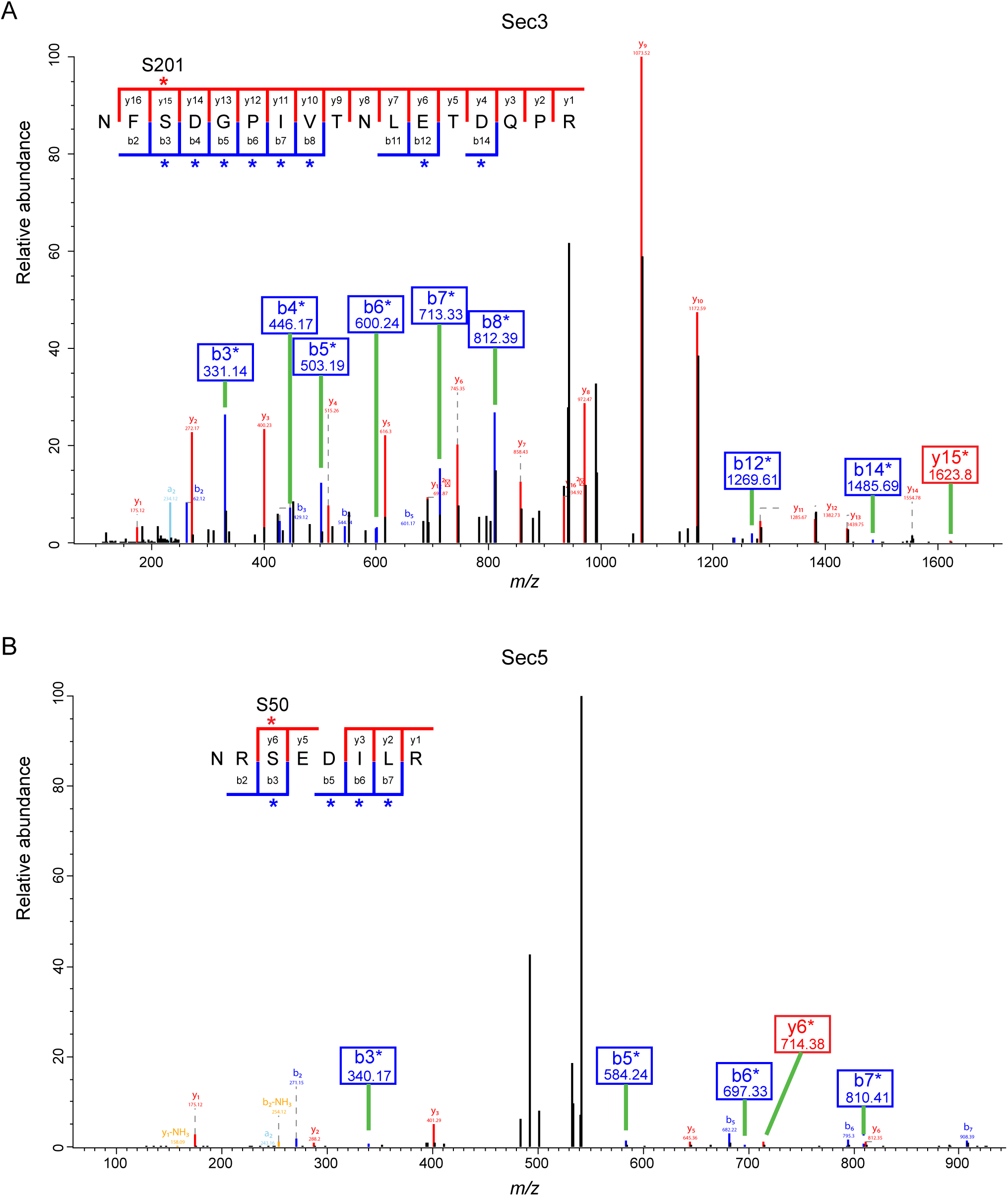
Orb6 phosphorylates recombinant GST-Sec3 and GST-Sec5 in vitro. **(A,B)** MS/MS spectra of phosphopeptides from kinase reactions shown in Figure 4D, confirming phosphorylation of Sec3 serine-201 (A) and Sec5 serine-50 (B). b and y ions marked with asterisks are diagnostic for the identification and localization of phosphorylation.

**Supplementary Figure 5.**
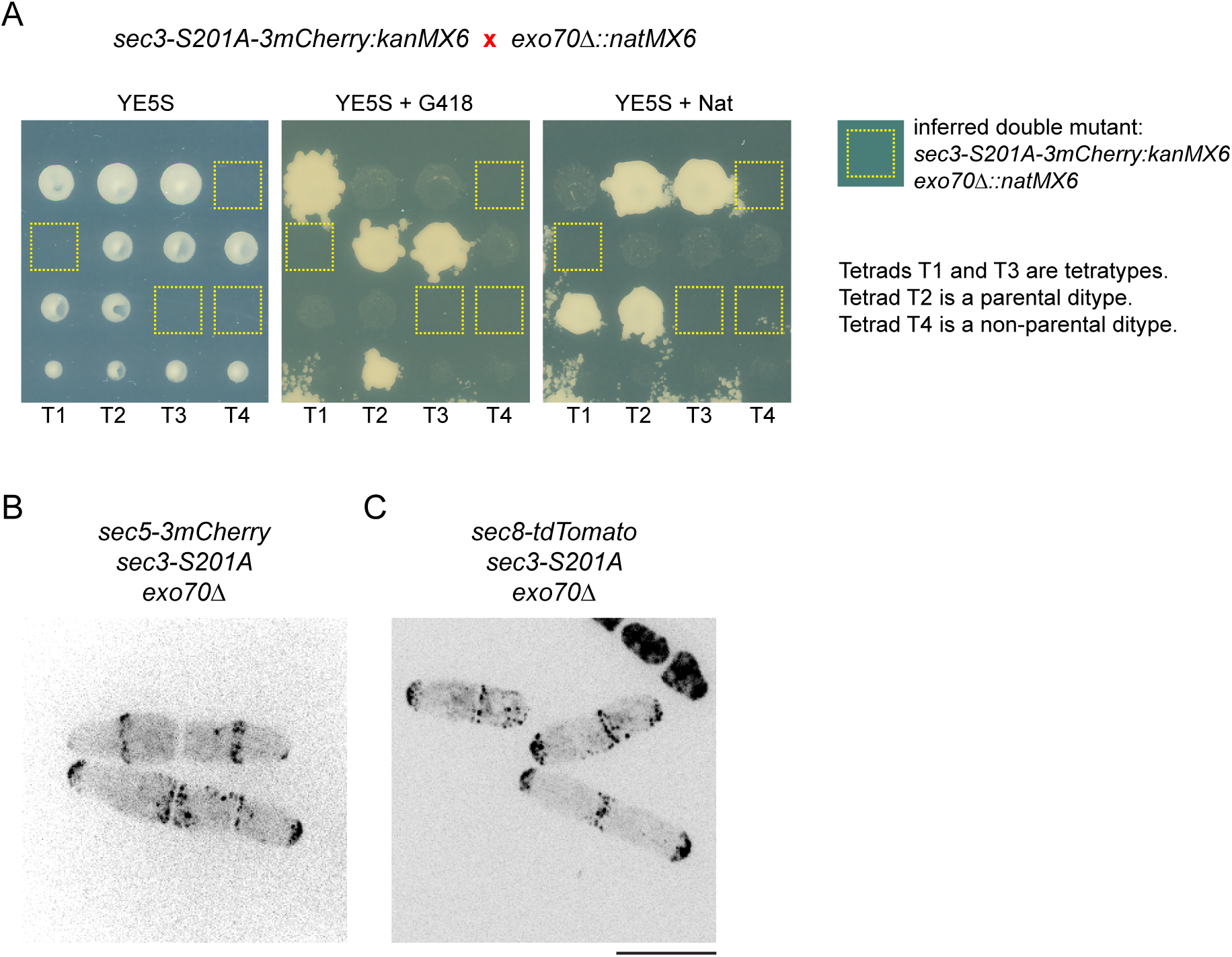
Localization of exocyst proteins in sec3-S201A exo70∆ cells. **(A)** *sec3-S201A-3mCherry* is synthetically lethal with *exo70∆.* As a result, Sec3-S201A-3mCherry localization in *exo70∆* background cannot be investigated. **(B, C)** Localization of Sec5-3mCherry (B) and Sec8-tdTomato (C) to cell tips and to the septation zone in *sec3-S201A exo70∆* backgrounds. In both strains, sister cells are joined in both cases, indicating a cell-separation defect (see also Fig. 6).

